# Metabolic reallocation in spinal cord oligodendrocytes drives chronic pain via neuronal β-amyloid production

**DOI:** 10.64898/2026.01.23.701389

**Authors:** Yannick Fotio, Saeed Al Masri, Zechuan Shi, Johnny Le, Sudeshna Das, Varvara I. Rubtsova, Alex Mabou Tagne, Cholsoon Jang, Vivek Swarup, Daniele Piomelli

**Affiliations:** Department of Anatomy and Neurobiology, University of California Irvine, Irvine, CA, USA; Department of Neurobiology and Behavior, University of California, Irvine, Irvine, CA, USA; Department of Biological Chemistry, University of California Irvine, Irvine, CA, USA; Department of Pharmaceutical Sciences, University of California Irvine, Irvine, CA, USA

**Author notes:** Lead contact: Daniele Piomelli. Gillespie Neuroscience Res. Facility (room 3101), 837 Health Sciences Rd., Irvine, CA 92697. Department of Biological Sciences, Center for Advances Pain Studies, The University of Texas at Dallas, Dallas, TX, USA.

## Abstract

Peripheral injury reprograms metabolism in spinal cord oligodendrocytes, initiating a molecular cascade that drives chronic pain via neuronal β-amyloid (Aβ) release. After injury, mouse spinal oligodendrocytes downregulate myelin protein synthesis and upregulate lipid biosynthesis—but reroute lipids toward neuroplastic remodeling and away from myelin maintenance. This metabolic reallocation disrupts myelin integrity and axonal function, causing neuronal accumulation of amyloid precursor protein, enhanced expression of its processing β-secretase BACE1, and local release of Aβ peptides. Blocking Aβ production or clearing Aβ deposits stops the transition to pain chronicity. Deleting the lysosomal lipid hydrolase NAAA in oligodendrocytes prevents both injury-induced Aβ production and chronic pain development. The findings identify an unexpected mechanistic link between chronic pain and Alzheimer’s-like neurodegeneration, positioning Aβ as a target for therapeutic intervention.

## Introduction

Pain affects millions worldwide and often persists long after healing is complete, evolving into a chronic debilitating condition that resists therapy (*1, 2*). This transition does not reflect partial recovery, but rather a fundamental reorganization of nociceptive processing, the molecular underpinnings of which remain unclear. Maladaptive plasticity in peripheral and central neurons has been implicated (*3–5*), alongside neuroimmune interactions involving microglia, astrocytes, and monocytes (*6–8*). Yet, these mechanisms alone do not fully explain how pain progresses to chronicity.

In mice, peripheral injury reprograms core metabolism in afferent segments of the spinal cord, causing a local decline in mitochondrial respiration (*9, 10*). This shift peaks four days after injury and coincides with upregulation of N-acylethanolamine acid amidase (NAAA) (*11*), a lysosomal hydrolase that degrades palmitoylethanolamide (PEA) and other endogenous ligands of the master regulator of energy and lipid homeostasis, peroxisome proliferator-activated receptor-α (PPAR-α) (*12, 13*). NAAA inhibition during this time window—but not before or after—prevents both metabolic reprogramming and the development of persistent pain across multiple models (*9*).

Here, we show that somatic injury uniquely alters metabolism in spinal oligodendroglia, impairing myelin upkeep and driving neuronal β-amyloid (Aβ) release and chronic pain development. Following injury, oligodendrocytes upregulate NAAA, downregulate myelin protein expression, and enhance phospholipid and sphingolipid biosynthesis—but redirect lipids toward neuroplastic remodeling at the expense of myelin maintenance. This metabolic reallocation disrupts myelin integrity and axonal function, causing neuronal accumulation of amyloid precursor protein (APP), upregulation of the APP-processing β-secretase BACE1, and production of Aβ peptides. These aggregate into plaque-like deposits, but only after pain has become persistent. Clearing Aβ or blocking its production post-injury prevents the transition to chronic pain. Moreover, NAAA deletion in oligodendrocytes identifies this enzyme as a control point of both neuronal Aβ production and pain chronification. These findings uncover a critical role for oligodendrocytes in the pathogenesis of chronic pain, identify an unexpected mechanistic link to Alzheimer’s-like neurodegeneration, and position Aβ as a potential therapeutic target.

## Results

### Peripheral injury reprograms metabolism in spinal oligodendrocytes

To identify cellular drivers of spinal metabolic reprogramming, we injected formalin (1%) in the hind paw of male mice and harvested ipsilateral lumbar (L4-L6) hemicords for single-nucleus RNA sequencing (snRNA-seq) four days later, when reprogramming peaks (*9, 10*) (Fig. 1a). Saline-injected mice served as controls. Though typically used to model acute pain, this paradigm reproduces key features of human chronic pain—including bilateral hypersensitivity and emotional, cognitive, and autonomic dysfunction (*9*). Following quality filtering, we analyzed 294,900 high-quality nuclei, recovering all major spinal cord cell types—including neurons, astrocytes, oligodendrocytes, and oligodendrocyte precursor cells (OPCs) (Fig. 1b). Cell-type composition remained largely stable, with only modest post-injury decreases in microglia and vascular leptomeningeal cells (Fig. S1a).

**Figure 1:**
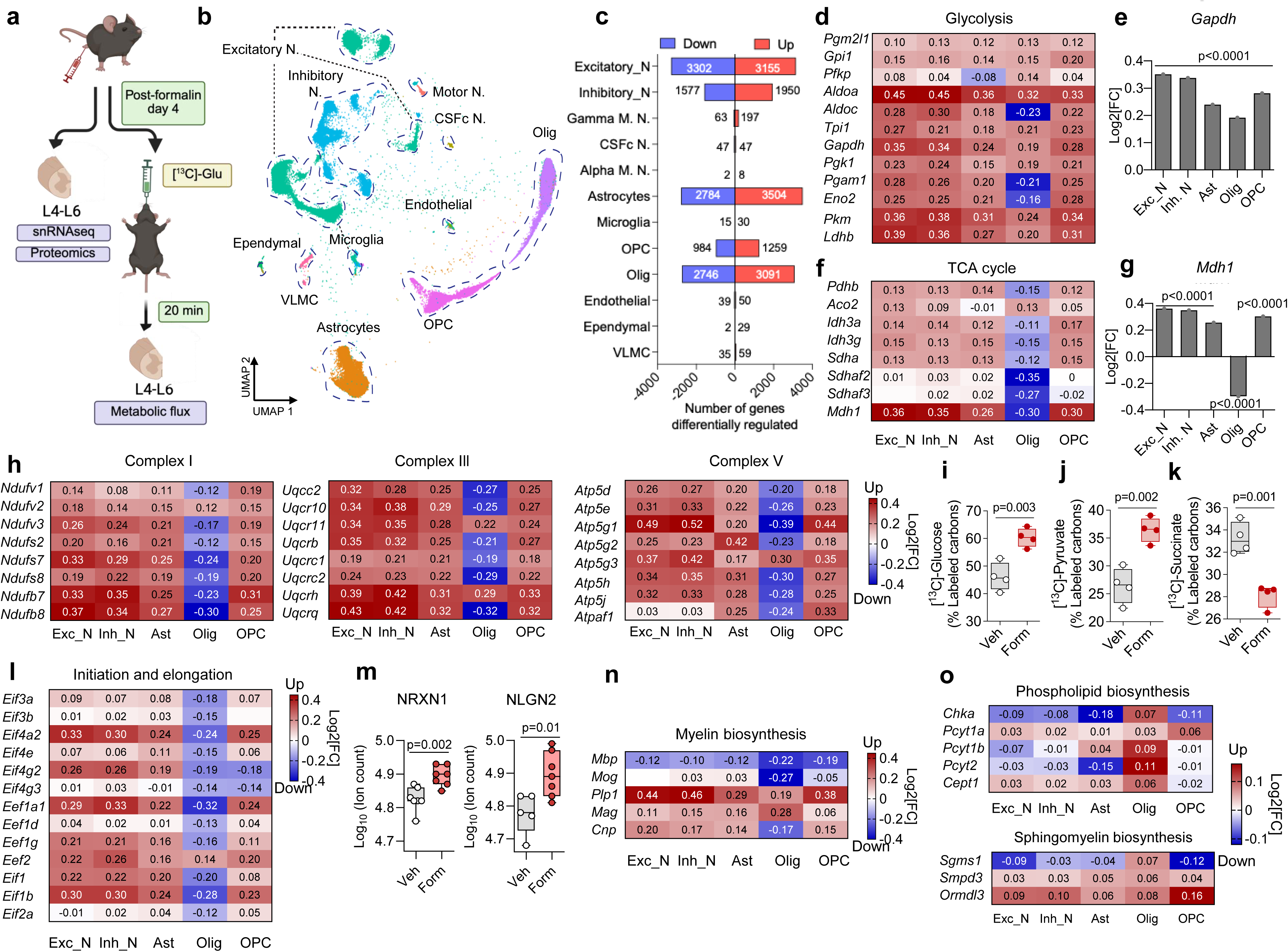
Peripheral injury reprograms core metabolism in spinal oligodendrocytes. (**a**) Male mice received intraplantar formalin (1%) or saline. After 4 days, ipsilateral lumbar (L4–L6) hemicords were collected for snRNA-seq and proteomics (n=4-7/group). A separate group received oral [¹³C]-glucose (10 mg/kg) 20 min before collection of L4–L6 hemicords for metabolic flux analysis. Image created with Biorender. (**b**) UMAP projection of cell types identified by snRNA-seq. Cell-type identities were assigned to clusters based on expression of canonical marker genes. (**c**) Number of differentially expressed genes detected in each cell-type cluster. Red, upregulated; blue, downregulated. (**d, f**) Heatmaps showing Log_2_ fold changes (formalin vs. vehicle) in expression of genes involved in glycolysis (**d**) and the tricarboxylic acid (TCA) cycle (**f**). (**e, g**) Log_2_ fold changes in expression of glycolytic enzyme glyceraldehyde-3-phosphate dehydrogenase (*Gapdh*) (**e**) and TCA enzyme malate dehydrogenase (*Mdh1*) (**g**). (**h**) Heatmaps showing Log_2_ fold changes in expression of genes encoding mitochondrial complexes I, III, and V. (**i–k**) Boxplots showing quantification of [¹³C]-carbon incorporation into glucose (**i**), pyruvate (**j**), and succinate (**k**). Data are expressed as percentage of total [¹³C]-labeled carbons. Gray, vehicle; red, formalin. (**l**) Log_2_ fold changes in expression of genes involved in the initiation and elongation of protein synthesis. (**m**) Proteomic quantification of neurexin 1 (NRXN1) and neuroligin 2 (NLGN2). Data are expressed as Log_10_ of ion counts. (**n, o**) Log_2_ fold changes in expression of genes involved in the biosynthesis of (**n**) myelin proteins and (**o**) phospholipids (top) and sphingolipids (bottom). Abbreviations: CSFc, cerebrospinal fluid-contacting neurons; Exc., excitatory; Inh. inhibitory; M., motor; N., neurons; Olig., oligodendrocytes; OPC, oligodendrocyte precursor cells; VLMC, vascular leptomeningeal cells. *Statistical analyses:* (**e, g**) Unpaired Student’s *t*-test with false discovery rate (FDR) correction. Bars represent mean Log_2_ fold changes of formalin vs. vehicle; (**i–k, m**) unpaired Student’s *t*-test. (**e, g**). Individual dots represent mice (n = 4-6 per group).

Differential gene expression analysis revealed widespread transcriptional changes across cell populations (Fig. 1c). Neurons, astrocytes, and OPCs showed coordinated upregulation of glycolytic genes, while oligodendrocytes exhibited a more heterogeneous response (Fig. 1d, e). Genes encoding tricarboxylic acid (TCA) cycle and oxidative phosphorylation components were upregulated in neurons, astrocytes, and OPCs, but were strongly downregulated in oligodendrocytes (Fig. 1f–h; Fig. S1b). These data suggest that—unlike other spinal cell types—oligodendrocytes suppress mitochondrial respiration following injury.

To assess metabolic consequences, we quantified glycolytic and TCA flux (*14*) combining oral [^13^C]-glucose administration with LC-MS analysis (Fig. 1a). Compared to vehicle controls, formalin-injected mice showed elevated [^13^C]-glucose in ipsilateral L4-L6 hemicords (Fig. 1i), consistent with increased uptake. This interpretation is supported by prior 2-deoxyglucose imaging studies (*15*) and the upregulation of glucose transporter *Slc2a1* (Glut1) (Fig. S2a). [^13^C]-labeling was also higher in glycolytic intermediates (Fig. S2b) and end-products, pyruvate (Fig. 1j) and lactate (Fig. S2b), but lower in succinate (Fig. 1k) and other TCA cycle intermediates (Fig. S2c), indicating reduced entry of glucose-derived carbons in the mitochondrial respiratory pathway. These changes are likely driven by the unique transcriptional reprogramming seen in oligodendrocytes, which comprise ∼30% of cells in the mouse CNS (*16*), although contributions from other cell populations cannot be excluded.

In injured mice, transcripts involved in the initiation and elongation of protein synthesis were higher in most spinal cell types, relative to uninjured controls, but sharply lower in oligodendrocytes (Fig. 1l). This coincided with local accumulation of various synaptic proteins, including neurexin 1 and neuroligin 2 (Fig. 1m; Fig. S3a, b)—identified by proteomics—and widespread transcriptional modifications (Fig. S4a, Data S1). Notably, oligodendrocytes reduced transcription of myelin-related genes (Fig. 1n, Fig S4b) but upregulated genes involved in phospholipid and sphingomyelin biosynthesis (Fig. 1o, Fig S4c, d), suggesting alterations in myelin composition and lipid homeostasis.

### Peripheral injury alters myelin composition in spinal cord

To assess whether myelin composition is affected, we performed proteomic and lipidomic analyses of myelin isolated on day 4 post-formalin from bilateral L4-L6 spinal segments (Fig. 2a). Principal component analysis (PCA) of proteomic data showed no separation between injured and non-injured groups (Fig. 2b), indicating limited myelin protein remodeling. Despite transcript downregulation (Fig. 1n), long-lived myelin components, myelin basic protein (MBP) and myelin oligodendrocyte glycoprotein (MOG), were only modestly and non-significantly reduced compared to controls (Fig. 2c, d). However, levels of the myelin-associated inhibitor oligodendrocyte-myelin glycoprotein (OMgp) (*17*) were significantly lower in injured relative to control mice (Fig. 2e), suggesting vulnerability of specialized myelin constituents at this early timepoint.

**Figure 2:**
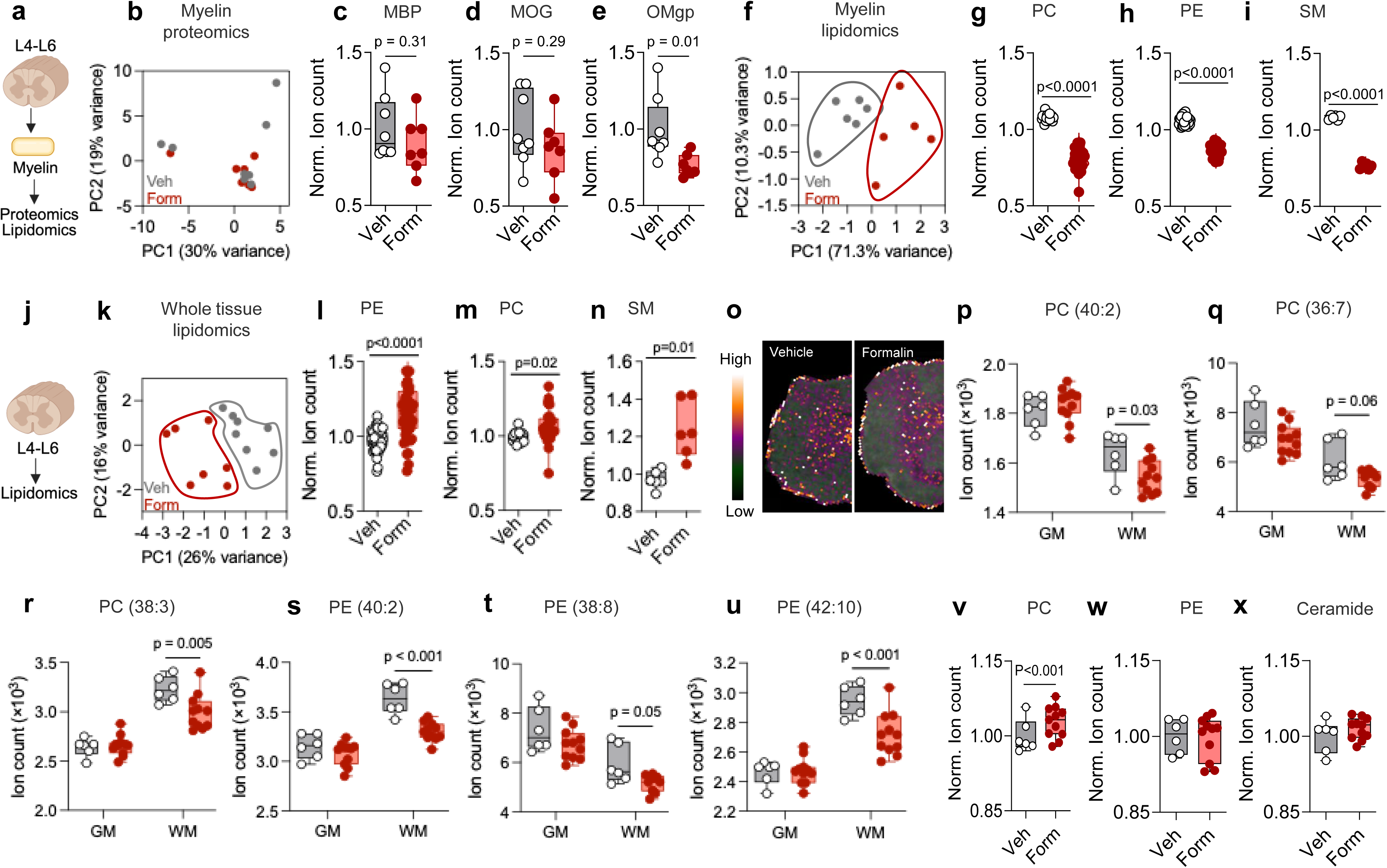
Peripheral injury alters myelin composition in spinal cord. (**a**) We isolated myelin from bilateral L4–L6 spinal cords 4 days after formalin or vehicle injection for proteomic or lipidomic analyses. (**b**) PCA of myelin proteomic data from mice treated with formalin (red symbols) or vehicle (gray symbols). (**c-e**) Boxplots showing levels of myelin basic protein (MBP, **c**), myelin oligodendrocyte glycoprotein (MOG, **d**), and oligodendrocyte myelin glycoprotein (OMgp, **e**) in mice treated with formalin or vehicle. Data are expressed as normalized ion counts. (**f**) PCA of myelin lipidomic data. (**g-i**) Levels of phosphatidylcholine (PC, **g**), phosphatidylethanolamine (PE, **h**), and sphingomyelin (SM, **i**) in mice treated with formalin or vehicle. Data are expressed as normalized ion counts summed across individual lipid species within each class. (**j**) We harvested bilateral L4-L6 spinal cords 4 days post-formalin or vehicle injection for lipidomic analysis. (**k**) PCA of the spinal lipidome. (**l-n**) Boxplots showing levels of PC (**l**), PE (**m**), and SM (**n**), presented as in **(g-i)**. (**o-x**) Imaging mass spectrometry of L4-L6 spinal cord at 4 days post-formalin. (**o**) Distribution of selected phospholipids in gray matter (GM) and white matter (WM). Reconstruction based on the following PC and PE species: PC (40:2), PC (36:7), PC (38:3), PE (40:2), PE (38:8), and PE (42:10). (**p-u**) Individual signal intensities for the PC and PE species shown in **(o)**. (**v-x**) Cumulative signal intensities for PC, PE, and ceramide. Images in **(a)** and **(j)** were created with Biorender. *Statistical analyses:* In panels (**c–e, g–i, l–n**), data were analyzed using unpaired *t*-tests with false discovery rate (FDR) adjustment; (**v-x**), data were analyzed using unpaired *t*-tests; (**p-u**), two-way ANOVA was used followed by Bonferroni *post hoc* test. Data in all panels are presented as box-and-whisker plots, where the median and interquartile range are shown. Individual data points represent mice (n = 4-8 per group).

In contrast to the relative stability of the myelin proteome, the myelin lipidome underwent pronounced changes post-injury. PCA revealed clear separation between formalin- and vehicle- injected groups (Fig. 2f), driven by losses in phosphatidylcholine (PC) (Fig. 2g), phosphatidylethanolamine (PE) (Fig. 2h), sphingomyelin (SM) (Fig. 2i), and cholesterol (Fig. S5a)—all essential for myelin structure and function (*18*). Other complex lipids and fatty acids were also significantly lower in myelin from injured compared to control mice (Fig. S5b–g). This loss was confined to white matter: in contrast to the marked lipid loss seen in myelin, lipidomic analyses of whole bilateral L4-L6 spinal tissue (Fig. 2j) revealed higher phospholipid and sphingolipid content, when these classes were considered as a whole (Fig. 2k–n; Fig. S6a-c). Imaging mass spectrometry at day 4 post-injury confirmed this segregation, showing broad lipid depletion in the white matter of formalin-treated mice. Composite images (Fig. 2o) illustrate the impact of injury on six representative phospholipids across white and gray matter (segmentation shown in Fig. S7), with corresponding signal intensities reported in Figure 2p-t. In the gray matter, levels of multiple PC species were increased (Fig. 2v) with no detectable change in PE and ceramide (Fig. 2w, x). These findings align with the upregulation of lipid-synthesizing enzymes in spinal oligodendroglia (Fig. 1o), suggesting that, following injury, oligodendrocytes reallocate lipids away from myelin and toward gray matter compartments undergoing neuroplastic remodeling (*3–5, 19*).

### Peripheral injury disrupts spinal myelin structure and function

The changes in myelin composition were accompanied by alterations in its structure and function. Electron microscopy analyses at day 4 post-injury showed preserved *g*-ratio and myelin thickness (Fig. 3a, b), but a strong trend toward reduced axonal diameter (*P* = 0.06, Fig. 3b) pointed to the early emergence of axon-myelin disruption. Consistent with this, ex vivo diffusion tensor imaging at day 14 revealed lower fractional anisotropy and higher mean diffusivity in L4 and L5 segments of formalin-injected mice, relative to controls (Fig. 3c, d; Fig. S8), indicative of compromised axonal integrity (*20*). To evaluate functional consequences, we quantified APP, a marker of axonal pathology (*21*). Injury elevated APP immunoreactivity in lumbar spinal cord neurons, reaching significance by day 21 (Fig. 3e, f). Notably, *App* transcription was also higher across L4-L6 cell types (Fig. 3g), likely reflecting ongoing neuroinflammation (*22*). These results indicate that somatic injury disrupts axonal function in afferent segments of the spinal cord, leading to neuronal APP accumulation and enhanced *App* transcription.

**Figure 3:**
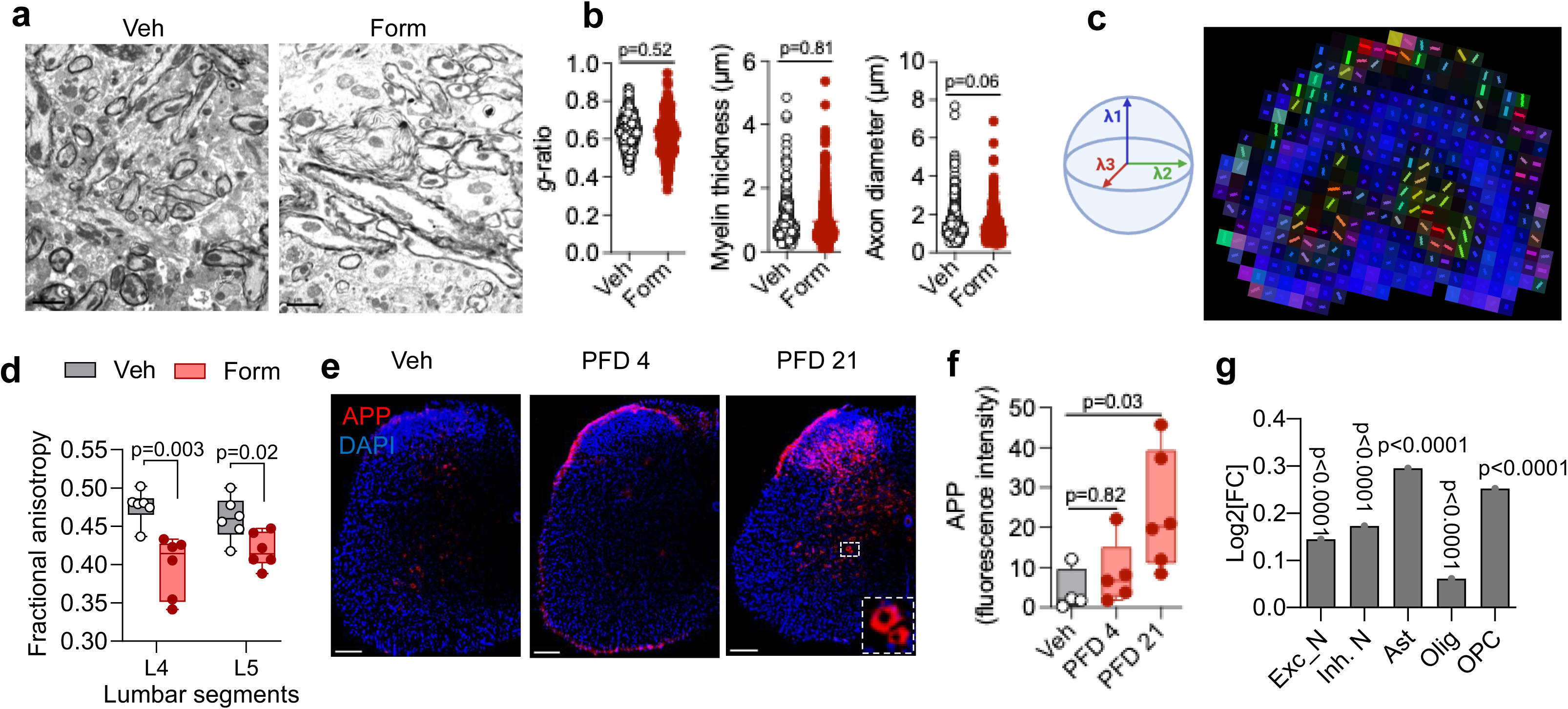
Peripheral injury disrupts spinal myelin structure and function. (**a**) Representative transmission electron microscopy images of lumbar (L4-L6) spinal cord sections 4 days post-injection of vehicle (left) or formalin (right). (**b**) *g*-ratio (left), myelin thickness (µm, middle), and axon diameter (µm, right) in mice treated with formalin (red symbols) or vehicle (gray symbols). Measurements are from 100 individual axons per mouse (n = 3 mice per group). (**c**) Diffusion tensor imaging (DTI). *Left*: Schematic of water diffusion modeled by three eigenvectors (λ₁, λ₂, λ₃) with magnitudes and principal orientations color-coded as blue (superior–inferior), red (anterior–posterior), and green (left–right). *Right*: Coronal reconstruction of the lumbar spinal cord from ex vivo DTI. (**d**) Boxplots showing fractional anisotropy in spinal segments L4 and L5 on day 14 after formalin or vehicle injection. (**e**) Fluorescent immunostaining of APP (red) in lumbar spinal sections at post-formalin days 4 and 21. Nuclei were stained with 4′,6-diamidino-2-phenylindole (DAPI, blue). Dashed boxes show APP-positive cells morphologically identifiable as neurons. Magnification: ×10 or zoom 4×. Scale bar, 100 μm. PFD, post-formalin day. (**f**) Immunoreactive APP levels (fluorescence intensity) in spinal cord of mice treated with formalin or vehicle. (**g**) Log₂ fold changes (formalin vs. vehicle) in *App* transcription across spinal cell types, assessed by snRNA-seq. *Statistical analyses:* (**b**) Nested Student’s *t*-test; (**d**) one-way ANOVA followed by Dunnett’s multiple comparison; (**f**) two-way ANOVA followed by Bonferroni *post hoc* test; (**g**) unpaired Student’s *t*-tests with false discovery rate (FDR) adjustment. Data in all panels are presented as box-and-whisker plots, where the median and interquartile range are shown. Individual data points represent mice, except in (**b**), where each dot represents an individual axon.

### Peripheral injury triggers spinal Aβ production

APP accumulation was accompanied by altered BACE1 expression. snRNA-seq of ipsilateral L4-L6 hemicords at day 4 post-injury showed modestly increased *Bace1* transcription in excitatory neurons (P = 0.05) and a marked decrease in oligodendrocytes (P < 0.001) (Fig. 4a). Mirroring these changes, immunoreactive BACE1 was elevated in neurons of formalin-compared to vehicle-injected mice (Fig. 4b; Fig. S9a, b). To assess whether Aβ is also produced, we performed sequential tissue extraction and found higher insoluble Aβ_42_ starting at day 4 post-formalin and persisting to day 21; Aβ_40_ was undetectable in this fraction (Fig. 4c). Soluble Aβ_40_ rose at day 21, with no detectable Aβ_42_ (Fig. 4d).

**Figure 4:**
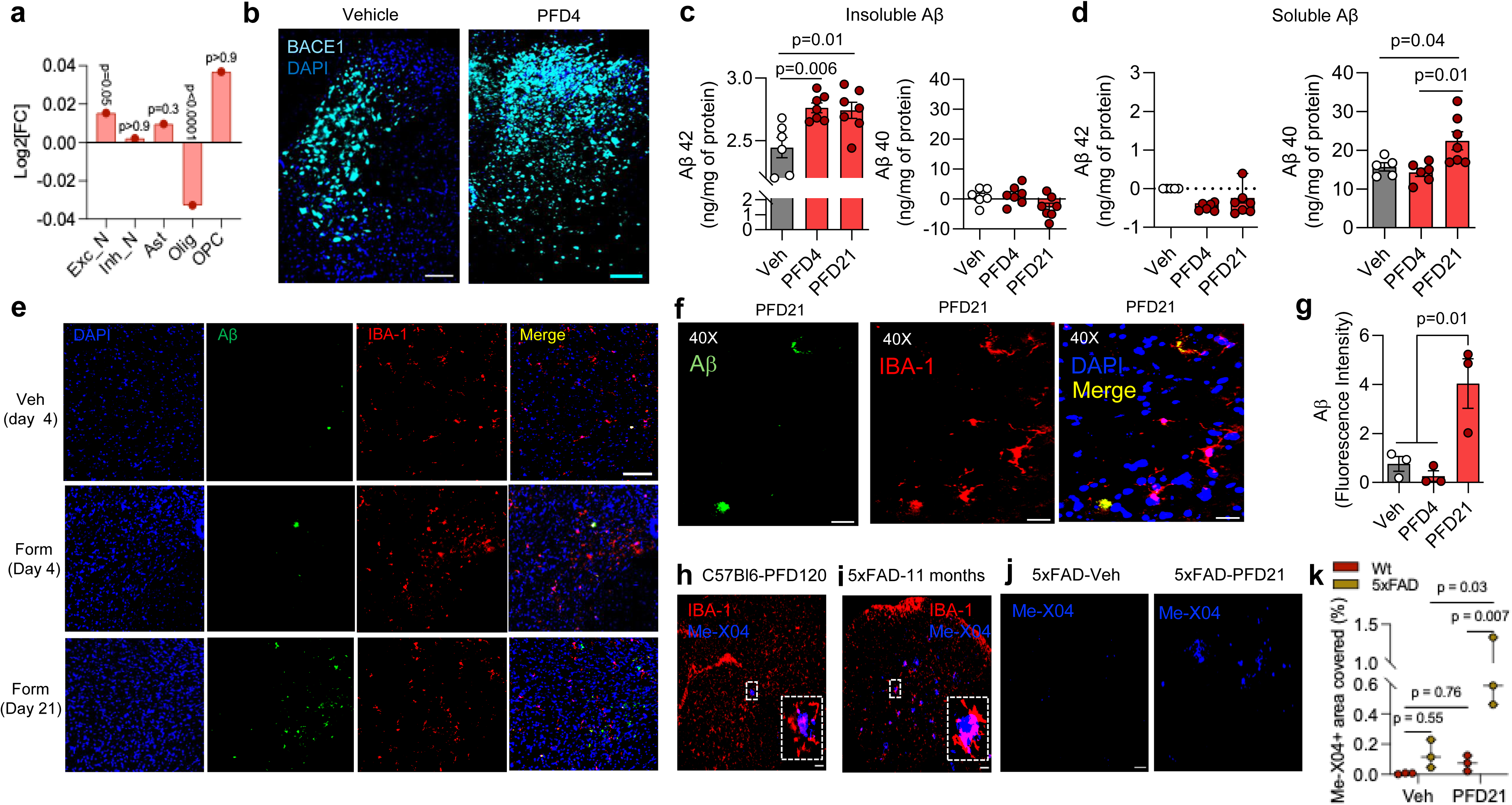
Peripheral injury triggers spinal Aβproduction. (**a**) Log_2_ fold changes (formalin vs. vehicle) in *Bace1* transcript across spinal cord cell types, assessed by snRNA-seq of ipsilateral lumbar (L4-L6) hemicords 4 days after injection of formalin or vehicle. Abbreviations: Exc., excitatory; Inh. inhibitory; N., neurons; Olig., oligodendrocytes; OPC, oligodendrocyte precursor cells. (**b**) Fluorescent immunostaining of BACE1 (cyan) in lumbar spinal sections 4 days after injection of vehicle (left) or formalin (right). Nuclei are stained with 4′,6-diamidino-2-phenylindole (DAPI, blue). Scale bar, 200 μm. PFD, post-formalin day. (**c, d**) Aβ_40_ and Aβ_42_ concentrations (ng/mg protein) in the insoluble (**c**) and soluble (**d**) fractions of L4-L6 spinal cords from mice treated with vehicle (gray symbols) or formalin (red symbols) at PFD4 and PFD21. (**e, f**) Fluorescent immunostaining of L4-L6 spinal sections showing Aβ deposits (green), IBA-1-positive microglia (red), and areas of colocalization (yellow) at PFD4 and PFD21. Nuclei are stained with DAPI. Magnification: ×10 (**e**) or ×40 (**f**); scale bar, 200 and 100 μm, respectively. (**g**) Immunofluorescence quantification of Aβ deposits in L4-L6 spinal sections of mice treated with formalin or vehicle at PFD4 and PFD21. (**h, i**) Representative images showing Me-X04–positive Aβ plaques (blue) and IBA1-positive microglia (red) in L4-L6 spinal sections from 8-month-old formalin-treated mice (PFD120) (**h**) and, for comparison, from 11-month-old naïve 5xFAD mice (**i**). Magnification: ×10 or ×40 (insets); scale bar, 100 μm. (**j**) Representative images showing Me-X04–positive plaques (blue) in 3-month-old 5xFAD mice 21 days after treatment with vehicle (left) or formalin (right). Magnification: ×10 or ×40 (insets); scale bar, 200 μm. (**k**) Immunofluorescence quantification of Me-X04–positive plaques in 3-month-old 5xFAD mice (yellow) or their wild-type littermates (red) 21 days after treatment with vehicle or formalin. *Statistical analyses:* (**a**) unpaired Student’s *t*-tests with false discovery rate (FDR) adjustment; (**c,d,k**) two-way ANOVA followed by Bonferroni *post hoc* test; (**g**) one-way ANOVA followed by Dunnett’s multiple comparison. Data are presented as box-and-whisker plots, where the median and interquartile range are shown. Individual data points represent mice, except in (**a**), where each dot represents mean Log_2_ fold changes of formalin vs. vehicle. Source data provided.

Longitudinal analysis revealed dynamic Aβ aggregation following injury. Aβ deposits were undetectable four days after formalin injection, but emerged by day 21 (Fig. 4e-g). Their frequent colocalization with IBA-1-positive microglia is consistent with early microglial clearance (*23*). At four months post-injury, methoxy-X04 staining (*24*) revealed mature fibrillar plaques in spinal cords of formalin- but not vehicle-treated mice (Fig. 4h; Fig. S10a, b). These deposits were morphologically indistinguishable from those seen in the 5×FAD model of familial Alzheimer’s disease (AD) (Fig. 4i; Fig. S10c) (*17*), and similarly surrounded by IBA-1-positive microglia (*25*). Although wild-type mice had fewer plaques than 5×FAD mutants, their presence indicates that peripheral injury can induce AD-like amyloid pathology in the spinal cord. Further supporting this, hind-paw formalin injection in 3-month-old 5×FAD mice—when plaques are normally undetectable—produced clear plaque accumulation, whereas vehicle had no such effect (Fig. 4j, k).

### Spinal Aβ release drives chronic pain

To assess whether Aβ release contributes to the development of chronic pain, we administered the monoclonal anti-Aβ antibody 4G8 (*26*) intrathecally once daily on days 2 and 4 after formalin (Fig. 5a). 4G8 treatment normalized insoluble Aβ_42_ and soluble Aβ_40_ levels (Fig. 5b) and suppressed contralateral heat and mechanical hypersensitivity, with effects on the latter emerging after one week (Fig. 5c; Fig. S11a). Ipsilateral hypersensitivity and paw edema were unaffected (Fig. S11b, c). The finding that 4G8 selectively prevents contralateral hypersensitivity, a prominent feature of this model (*9*), is consistent with a role for Aβ in persistent rather than acute nociception (*27, 28*). Confirming a link between spinal Aβ_42_ concentrations and pain chronification, Aβ_42_ levels were inversely correlated with heat withdrawal latency (R²=0.2; P<0.03) and mechanical withdrawal threshold (R²=0.3; P<0.004), measured on day 14 post-injury (Fig. 5d). 4G8 also prevented formalin-induced anxiety-like behavior (Fig. S11d), but not cognitive impairment (Fig. S11e)—two additional features of this model (*9*) and chronic pain in humans (*29*). The findings indicate that Aβ release contributes to the transition to pain chronicity following somatic injury.

**Figure 5.**
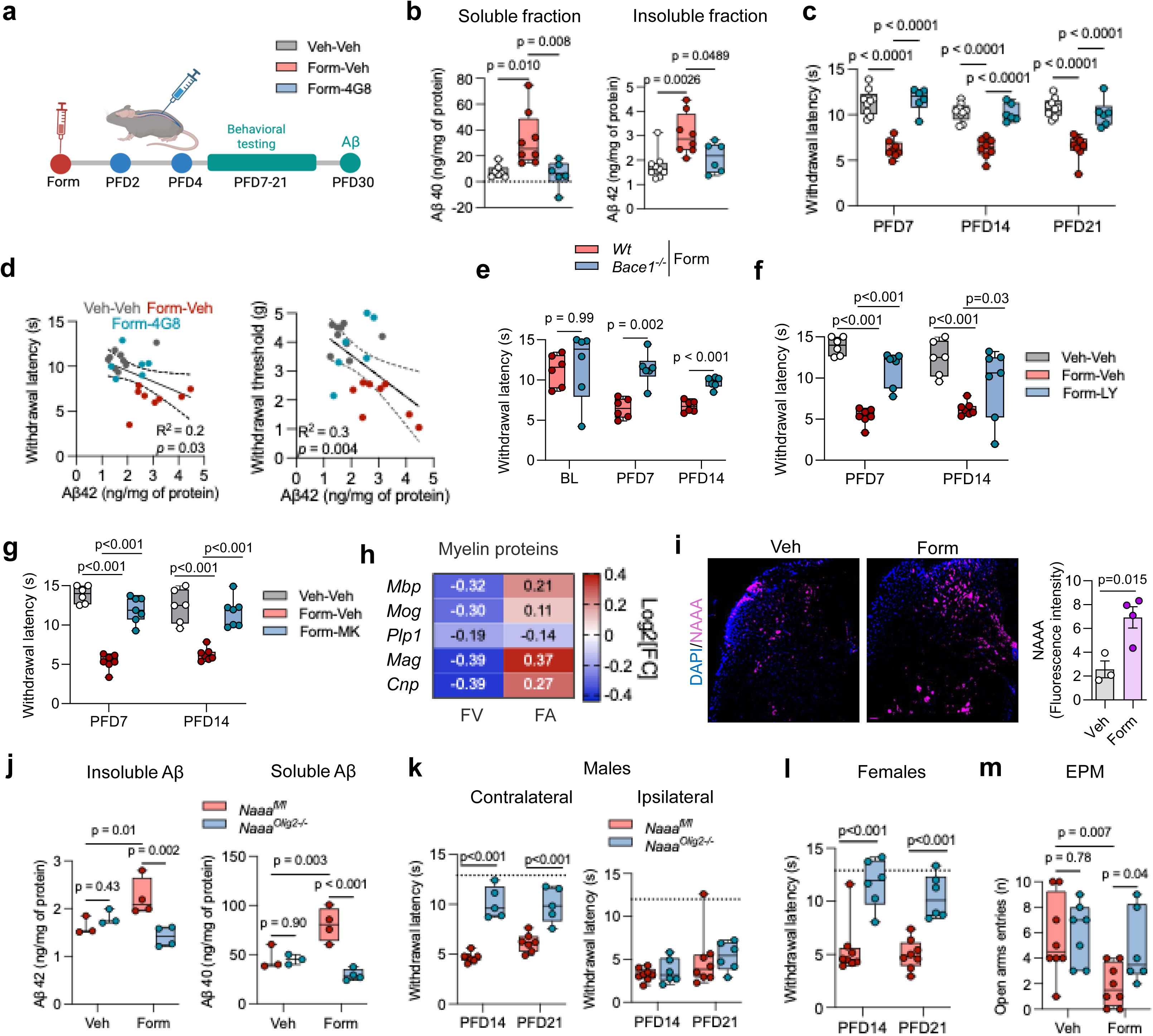
Spinal Aβ production drives chronic pain under control of oligodendrocyte NAAA. (**a**) Male mice received intrathecal injections of the Aβ-specific antibody 4G8 (1μg/μl, 10 μl) on post-formalin day (PFD) 2 and 4 (n = 6–9 per group). Mechanical and heat hypersensitivity were monitored over 21 days; mice were euthanized on PFD30 for Aβ quantification in lumbar (L4–L6) spinal cords. Image created with Biorender. (**b**) Boxplots showing Aβ_40_ and Aβ_42_ levels (ng/mg protein) in insoluble (left) and soluble (right) fractions. Symbols: vehicle-vehicle (gray), formalin-vehicle (red), and formalin-4G8 (blue). (**c**) Effects of 4G8 administration on contralateral heat hypersensitivity (latency, seconds) at PFD7, PFD14, and PFD21. (**d**) Correlation between Aβ_42_ levels at PFD21 and heat withdrawal latency (left; R² = 0.20, P = 0.03) or mechanical withdrawal threshold (right; R² = 0.30, P = 0.004). (**e**) Contralateral heat hypersensitivity in male *Bace1^-/-^*mice and wild-type littermates before formalin injection (baseline, BL) and at PFD7 and PFD14. Symbols: formalin-*Bace1^-/-^* mice (blue), and formalin-wild type (red). (**f, g**) Contralateral heat hypersensitivity at PFD7 and PFD14 in male wild-type mice treated with BACE1 inhibitors LY2811376 (10 mg/kg, IP) (**f**) or MK-8931 (10 mg/kg, IP) (**g**). (**h**) Effect of NAAA inhibitor ARN19702 (30 mg/kg, IP) administered at PFD2, 3, and 4 in male wild-type mice on myelin-related gene expression, assessed by bulk RNA-seq at PFD4. Three group of mice (n = 5 per group) were used: vehicle/vehicle; formalin/vehicle, and formalin/ARN19702. Values represent Log2 fold changes in gene expression comparing formalin-vehicle vs vehicle-vehicle (FV) or formalin-ARN19702 vs formalin-vehicle (FA). Red, upregulated; blue, downregulated. (**i**) Immunoreactive NAAA in male wild-type mice 4 days after injection of vehicle or formalin; right, quantification of fluorescent intensity. (**j**) Aβ_40_ and Aβ_42_ concentrations in insoluble and soluble fractions from formalin- or vehicle-treated *Naaa^Olig2−/−^*and *Naaa^fl/fl^* littermates. (**k**) Effects of oligodendrocyte-specific *Naaa* deletion on contralateral (left) and ipsilateral (right) heat hypersensitivity in male mice at PFD14 and PFD21. (**l**) Effects of oligodendrocyte-specific *Naaa* deletion on contralateral heat hypersensitivity in female mice at PFD14 and PFD21. (**m**) Anxiety-like behavior (number of open arm entries in an elevated plus maze) in formalin- and vehicle-treated *Naaa^Olig2−/−^* and *Naaa^fl/fl^* male littermates at PFD14. *Statistical analyses:* In panels (**b**), data were analyzed using one-way ANOVA followed by Dunnett’s *post hoc* test; (**c, e, f, g, j, k, l, m**), two-way ANOVA was used followed by Bonferroni *post hoc* test; (**i**) unpaired Student’s *t*-test. Data in all panels are presented as box-and-whisker plots, where the median and interquartile range are shown, except (**i**) that presents mean ± SEM. Individual data points represent mice. *P* values reflect comparisons between formalin-and vehicle-treated groups.

Additional support for this hypothesis came from genetic and pharmacological interventions targeting Aβ production. Unlike their wild-type littermates, mice constitutively lacking *Bace1* failed to develop contralateral heat hypersensitivity post-formalin (Fig. 5e; Fig. S12a-c). Furthermore, hypersensitivity was prevented by systemic administration on days 2-4 post-injury of either of two BACE1 inhibitors—LY2811376 (*30*) and MK8931 (*31*) (Fig. 5f, g)—or the γ-secretase inhibitor NGP555 (*32*) (Fig. S12d, e).

### NAAA in oligodendrocytes controls Aβ production and pain chronification

NAAA inhibition on days 2-4 post-injury blocks the progression to chronic pain across multiple models (*9*). The same intervention—using the inhibitor ARN19702 (30 mg/kg, IP) (*33*)—restores transcriptional programs linked to myelin biosynthesis (Fig. 5h) and mitochondrial respiration (Fig. S13a, b), as revealed by bulk RNA-seq of ipsilateral L4-L6 hemicords. Moreover, immunoreactive NAAA is elevated in spinal oligodendrocytes on day 4 post-injury, compared to controls (Fig. 5i, Fig. S13c).

To assess NAAA’s contribution to Aβ production, we generated mice selectively lacking *Naaa* in oligodendrocytes (*Naaa*^Olig2-/-^) (Fig. S14a). In male *Naaa*^Olig2-/-^ mutants, formalin injection did not increase spinal Aβ_42_ and Aβ_40_ levels at day 30, while increasing them in floxed littermates (Fig. 5j). Furthermore, male and female *Naaa*^Olig2-/-^ mice did not develop contralateral heat hypersensitivity (Fig. 5k and 5l) or anxiety-like behavior post-injury (Fig. 5m; Fig. S14b). Cognitive impairment was not affected (Fig. S14c). Lastly, *Naaa* deletion in oligodendrocytes decreased spinal Aβ_42_ and Aβ_40_ levels and normalized heat and mechanical sensitivity in the chronic constriction injury model of persistent pain (*34*) (Fig. S15c-c). We conclude that oligodendrocyte NAAA is a necessary upstream driver of neuronal Aβ release and the transition to pain chronicity in both sexes and in two models of somatic injury.

## Discussion

Our findings show that, following peripheral injury, spinal oligodendrocytes reduce myelin protein synthesis and reroute newly synthesized lipids from white to gray matter regions undergoing neuroplastic remodeling linked to central sensitization (*3–5, 19*). This metabolic reallocation destabilizes myelin, impairs axonal function, and leads to neuronal APP accumulation and Aβ release—events that we found are required for the transition to pain chronicity. The lipid hydrolase NAAA governs this cascade: its selective deletion in oligodendrocytes prevent Aβ buildup and halts pain chronification, providing a mechanistic basis for the proposed involvement of oligodendroglia in this process (*35*).

Unlike other spinal cell populations, which upregulate both glycolysis and mitochondrial respiration following somatic injury, oligodendrocytes adopt a distinct transcriptional program that suppresses TCA cycle activity, oxidative metabolism, and myelin protein synthesis while upregulating phospholipid and sphingolipid biosynthesis. This metabolic shift has two main consequences. First, it increases the availability of pyruvate and lactate, energy-rich glycolytic end-products that can be transferred to axons via monocarboxylate transporters (*36*) to support the hyperexcitability characteristic of central sensitization (*37*). Second, and key to explain our results, it increases lipid levels in the spinal gray matter while reducing them in white matter, suggesting a redistribution of these membrane components toward regions undergoing synaptic reorganization, potentially via vesicular or non-vesicular transport mechanisms (*38–40*). The loss of myelin lipids may be exacerbated by enhanced oligodendroglial fatty acid oxidation to sustain axonal function (*40*). This injury-induced reprogramming differs from other forms of oligodendrocyte plasticity, including adaptive myelination driven by neuronal activity in OPCs (*41*) and the transcriptional reconfiguration seen in late-stage AD pathology (*42*). Instead, it emerges in mature oligodendrocytes and disrupts myelin integrity within days of injury.

These alterations in myelin integrity create a permissive environment for Aβ production, reminiscent of that observed in AD models, where myelin dysfunction promotes Aβ release through APP accumulation and BACE1 induction (*24*). We found that peripheral injury exerts a similar—albeit much faster—effect in the spinal cord of otherwise healthy mice. Aβ release is detectable by day 4 post-formalin and is necessary for the transition to chronic pain. Immunological, genetic, and pharmacological interventions that block Aβ buildup in the days following injury also prevent the development of formalin-induced contralateral hypersensitivity and anxiety-like behavior—two defining features of chronic pain in this model (*9*)—without affecting ipsilateral responses, indicating that Aβ contributes to persistent but not acute nociception. These interventions also fail to rescue cognitive deficits post-formalin, consistent with evidence that the sensory, affective, and cognitive dimensions of chronic pain recruit distinct neural mechanisms (*29*). Notably, amyloid plaques surrounded by reactive microglia—hallmarks of AD pathology (*25*)—appear in the spinal cord only after chronic pain is established, suggesting that are unlikely to trigger its onset. The role of these plaques and of continued Aβ release in maintaining chronic pain are important questions for future investigation.

The signals that initiate oligodendroglial reprogramming are still undefined, but we identified the lysosomal lipid amidase NAAA as a critical checkpoint. Inhibiting NAAA 2-4 days after injury restores transcriptional programs governing myelin and lipid biosynthesis, while oligodendrocyte-specific *Naaa* deletion blocks Aβ production along with contralateral hypersensitivity and anxiety-like behavior. Notably, global *Naaa* deletion or systemic NAAA inhibition abrogates all molecular and behavioral hallmarks of chronic pain, suggesting a broader role for NAAA across multiple cell types. Although the downstream mechanism remains uncertain, one plausible model—supported by prior work (*9–11*)—is that NAAA reduces tonic activation of PPAR-α by degrading its endogenous agonist, PEA. This would not only impair PPAR-α–mediated stimulation of mitochondrial respiration and lipid oxidation (*43*), but also promote neuroinflammation (*44*) and BACE1 expression (*45*).

In humans, chronic pain is an independent risk factor for cognitive impairment and AD (*46–48*). Notably, in one cohort of 887 older men without dementia, persistent pain was associated with elevated plasma Aβ_40_ levels (*48*), echoing our findings in mice. While not conclusive, this clinical association reinforces the potential relevance of our results. Our data further suggest that early myelin dysfunction may forecast the progression to chronic pain, consistent with imaging studies linking white matter abnormalities to pain persistence (*49*). Together, these findings identify Aβ as a candidate therapeutic target for modifying the course of chronic pain following somatic injury.

## Supporting information

Raw data

## Acknowledgements

This project was funded by NIH grants 1R01DA055578-01 and 4R01AG065329 - 02 (to D.P.) as well as 1K99AT012658-01 (to Y.F.) and R01-AA029124 (to C.J.). We thank the University of California Irvine’s Mass Spectrometry Facility and Dr. Felix Grün for assistance with imaging mass spectrometry analyses, which were conducted on a Shimadzu iMScope 9050 funded by NIH grant S10OD034422-01. We also thank the University of California Irvine’s Department of Pathology’s Experimental Tissue Resources for support with electron microscopy and Dr. Andre Obenaus for help with diffusion tensor imaging.

## Data and materials availability

All data necessary to support the conclusion of this study are available in the manuscript or the supplementary information. Raw data are deposited in figshare repository (DOI: 10.6084/m9.figshare.29565233).

---

**Figure S1.**
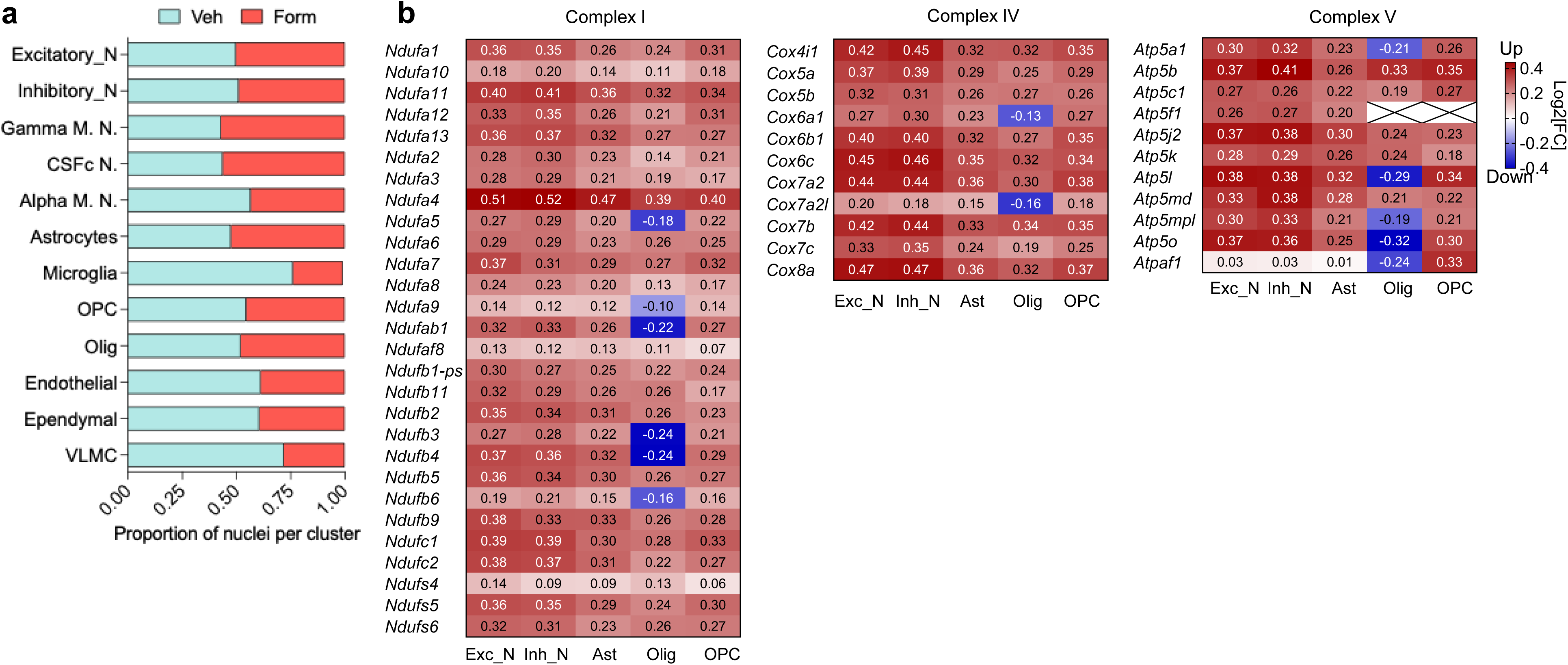

**Figure S2.**
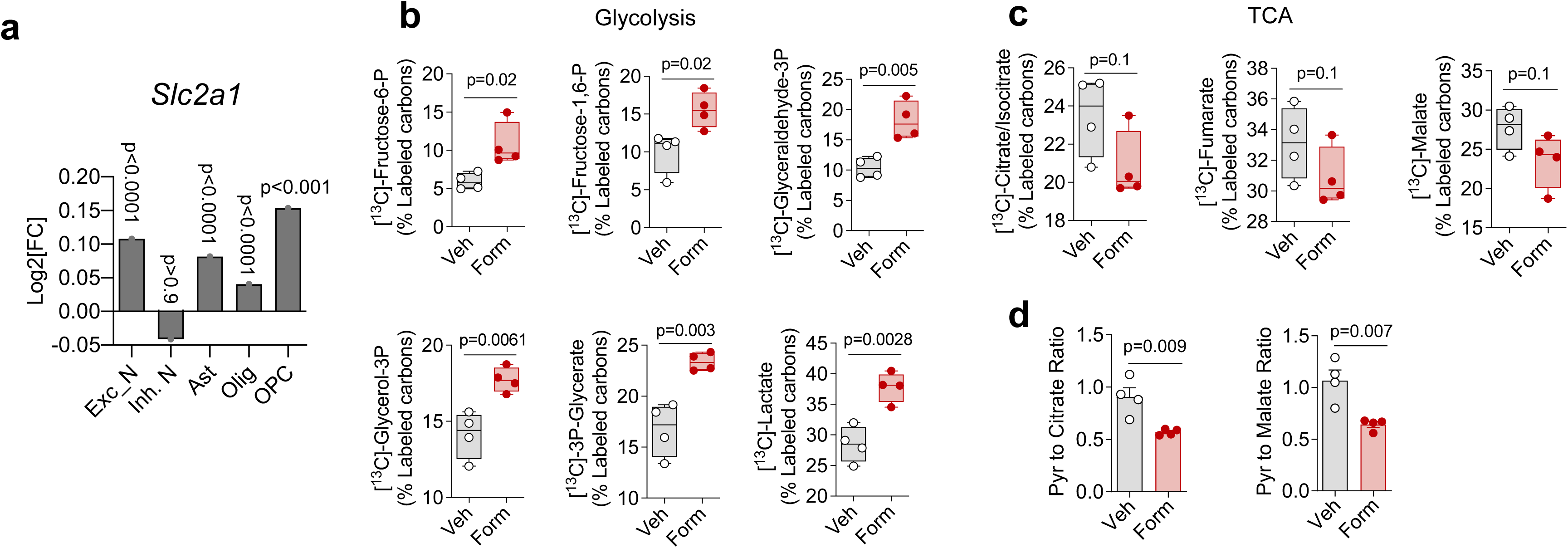

**Figure S3.**
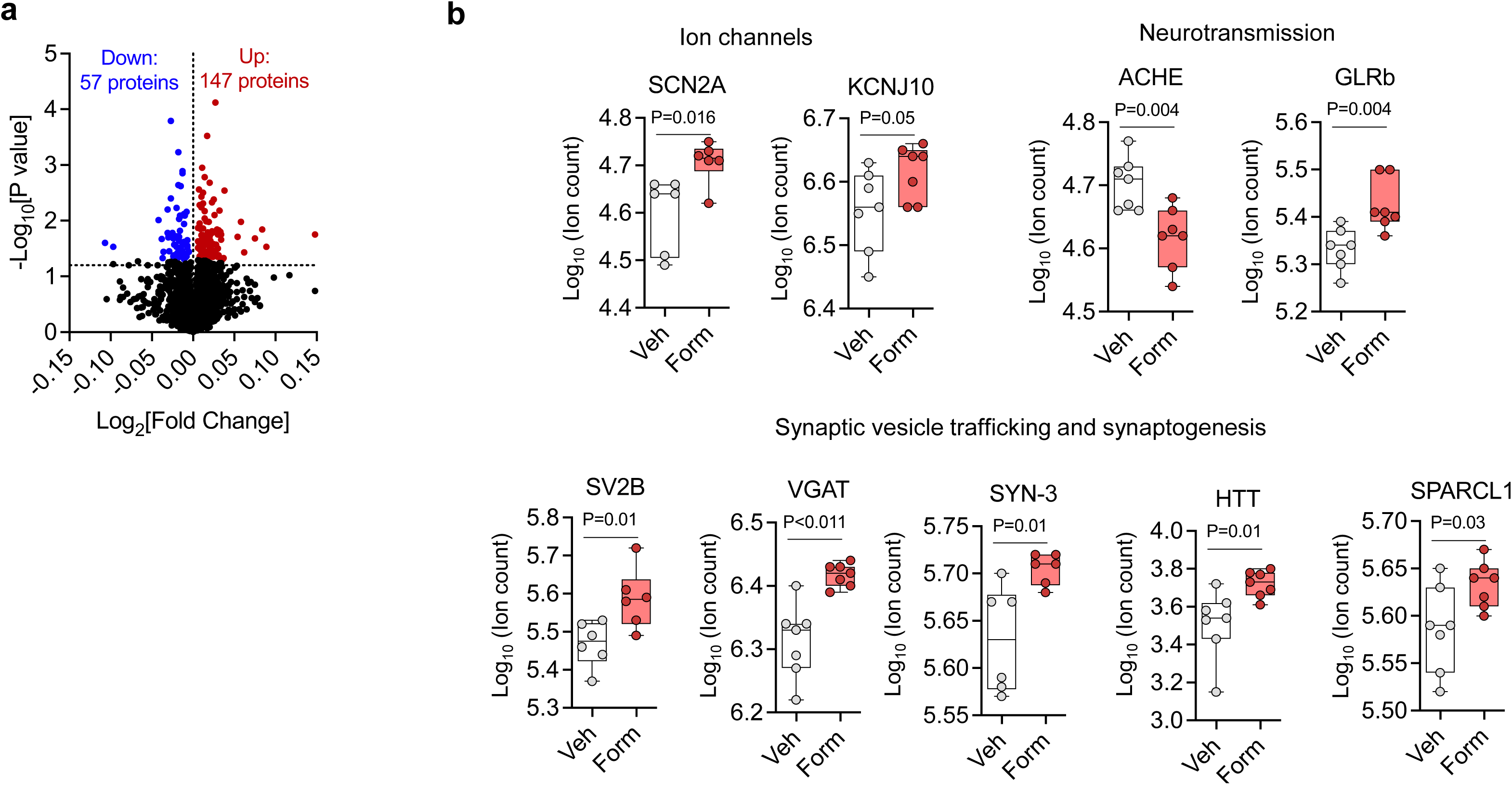

**Figure S4.**
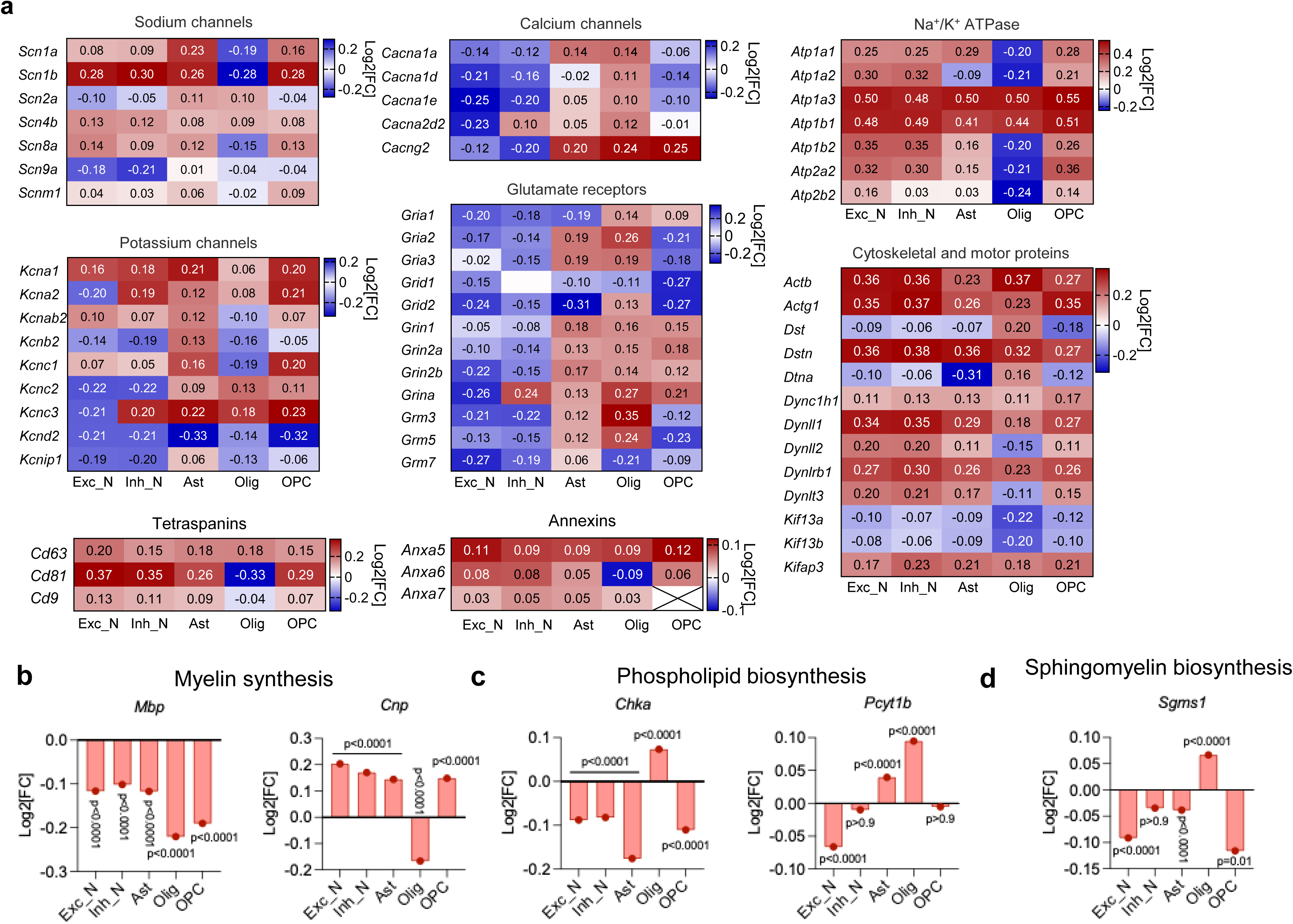

**Figure S5.**
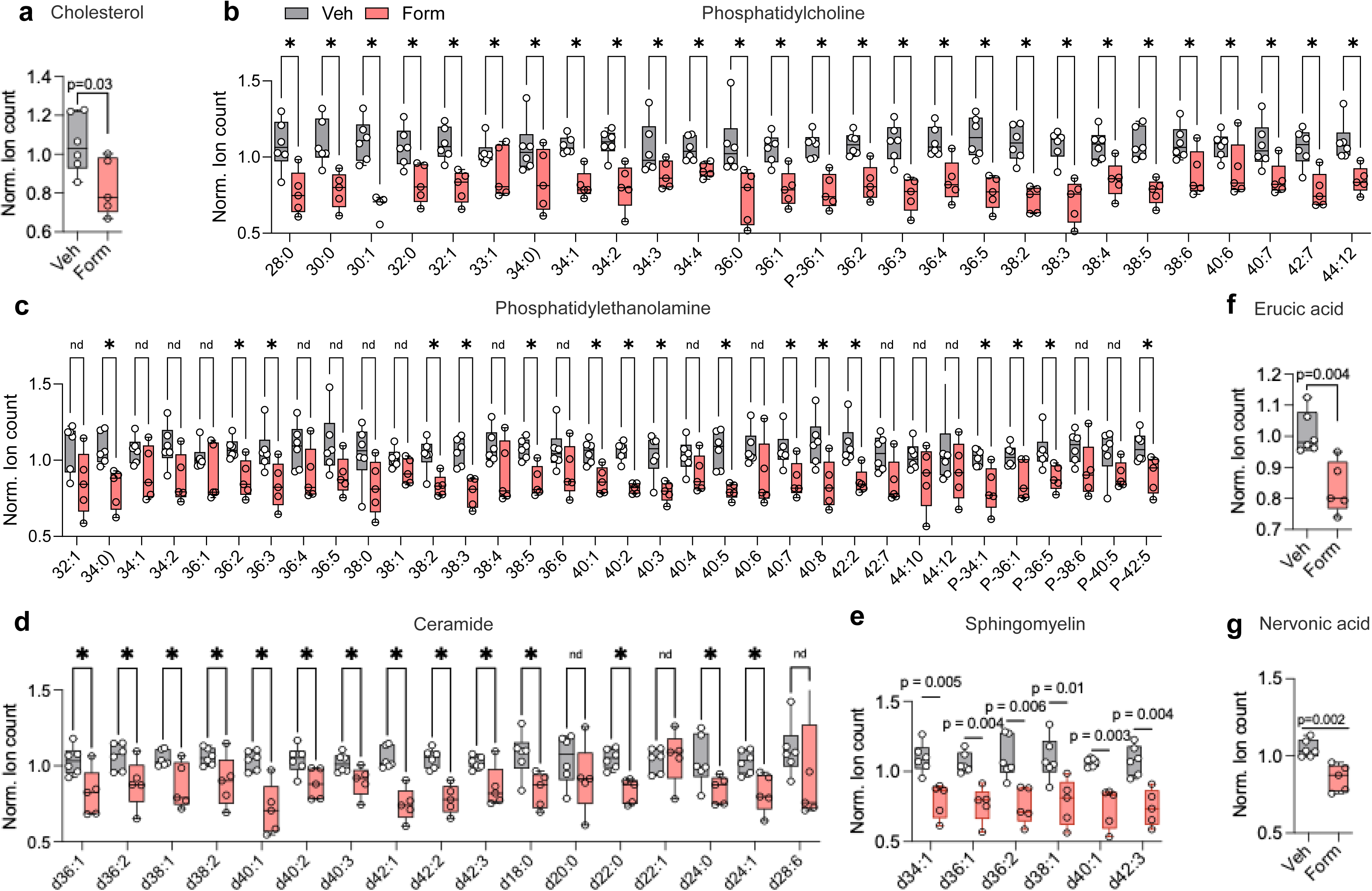

**Figure S6.**
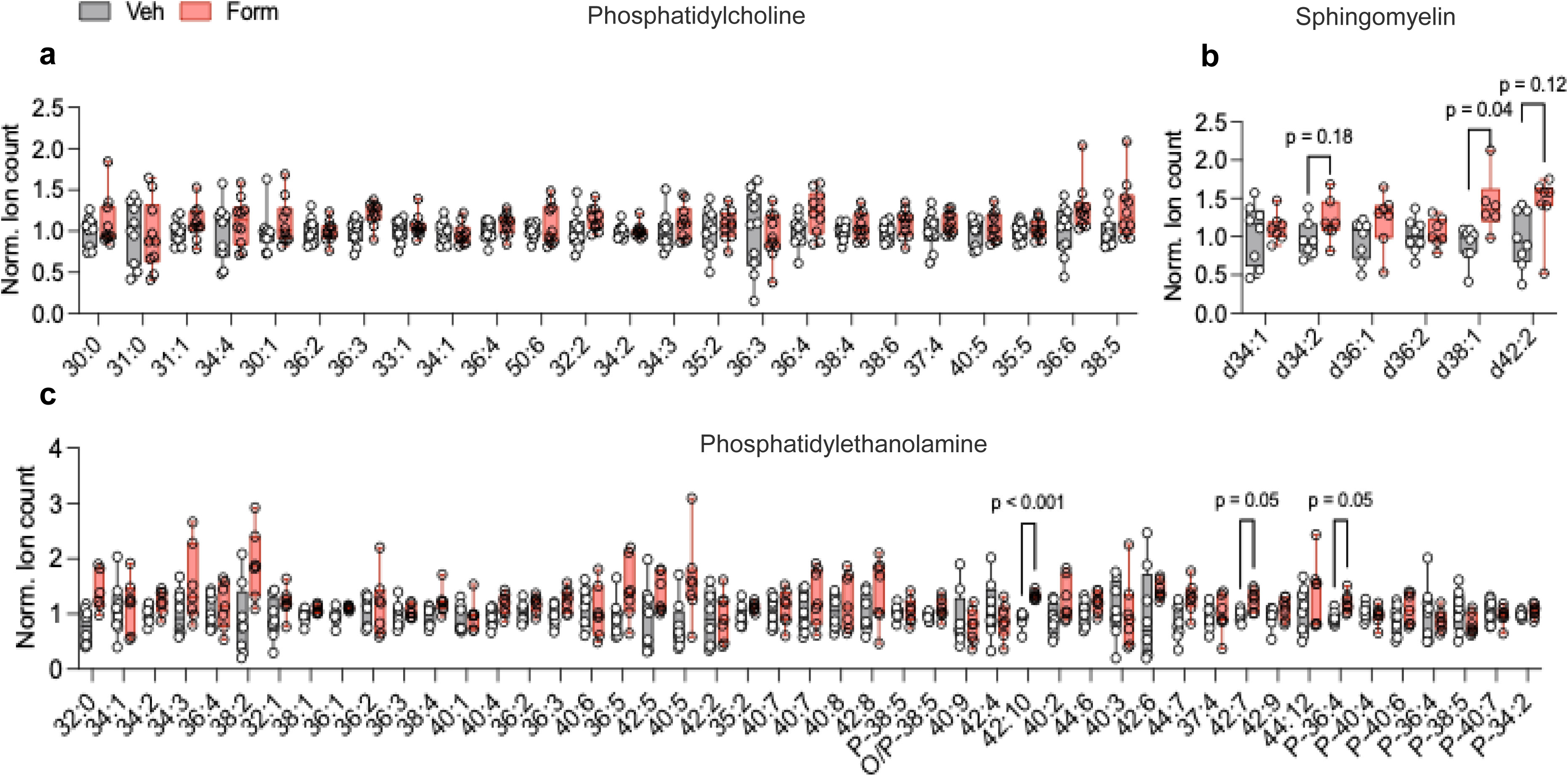

**Figure S7.**
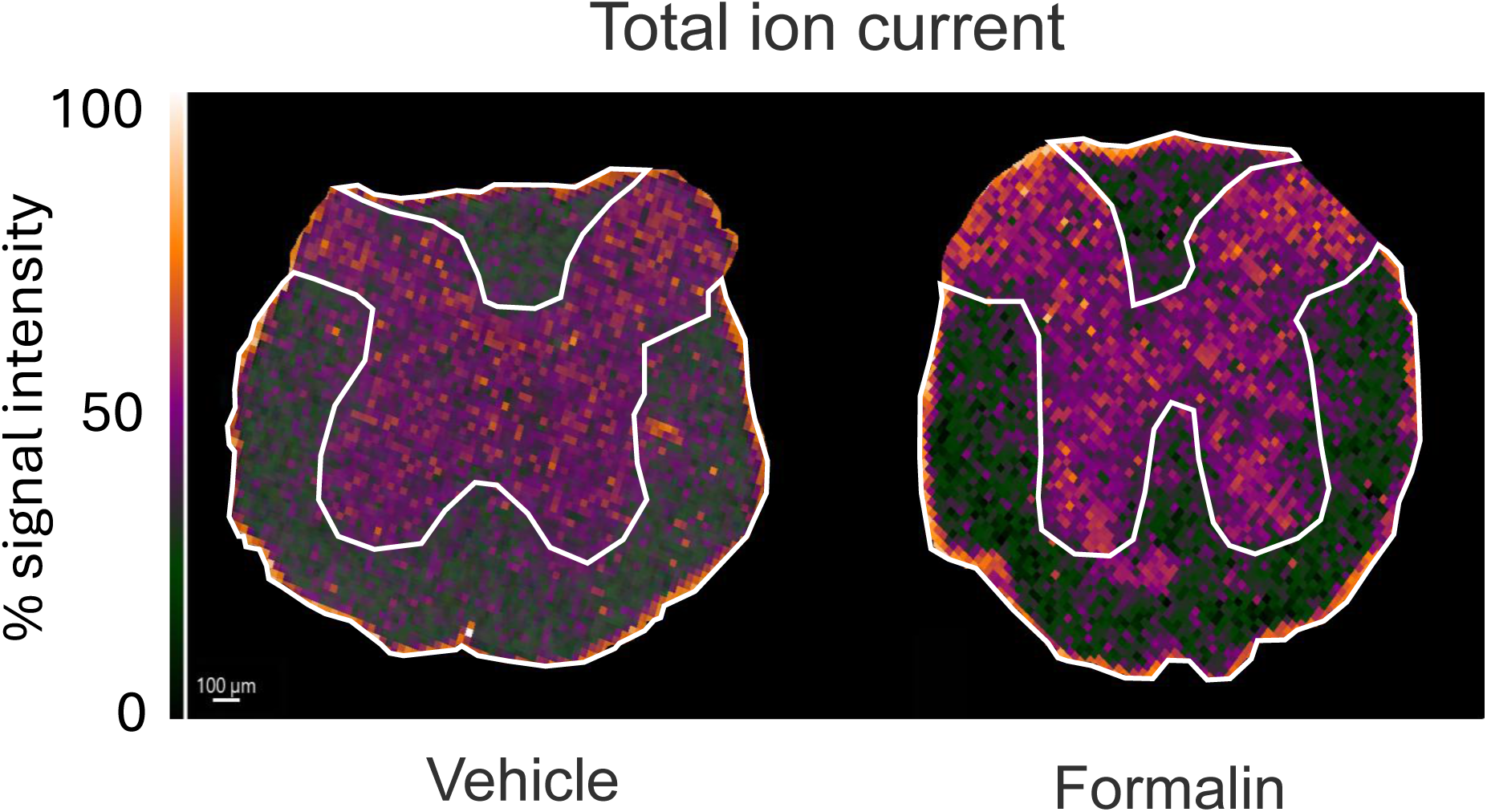

**Figure S8.**
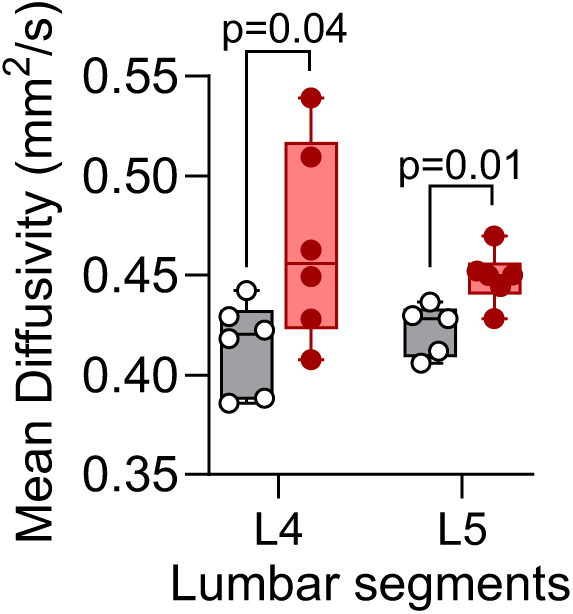

**Figure S9.**
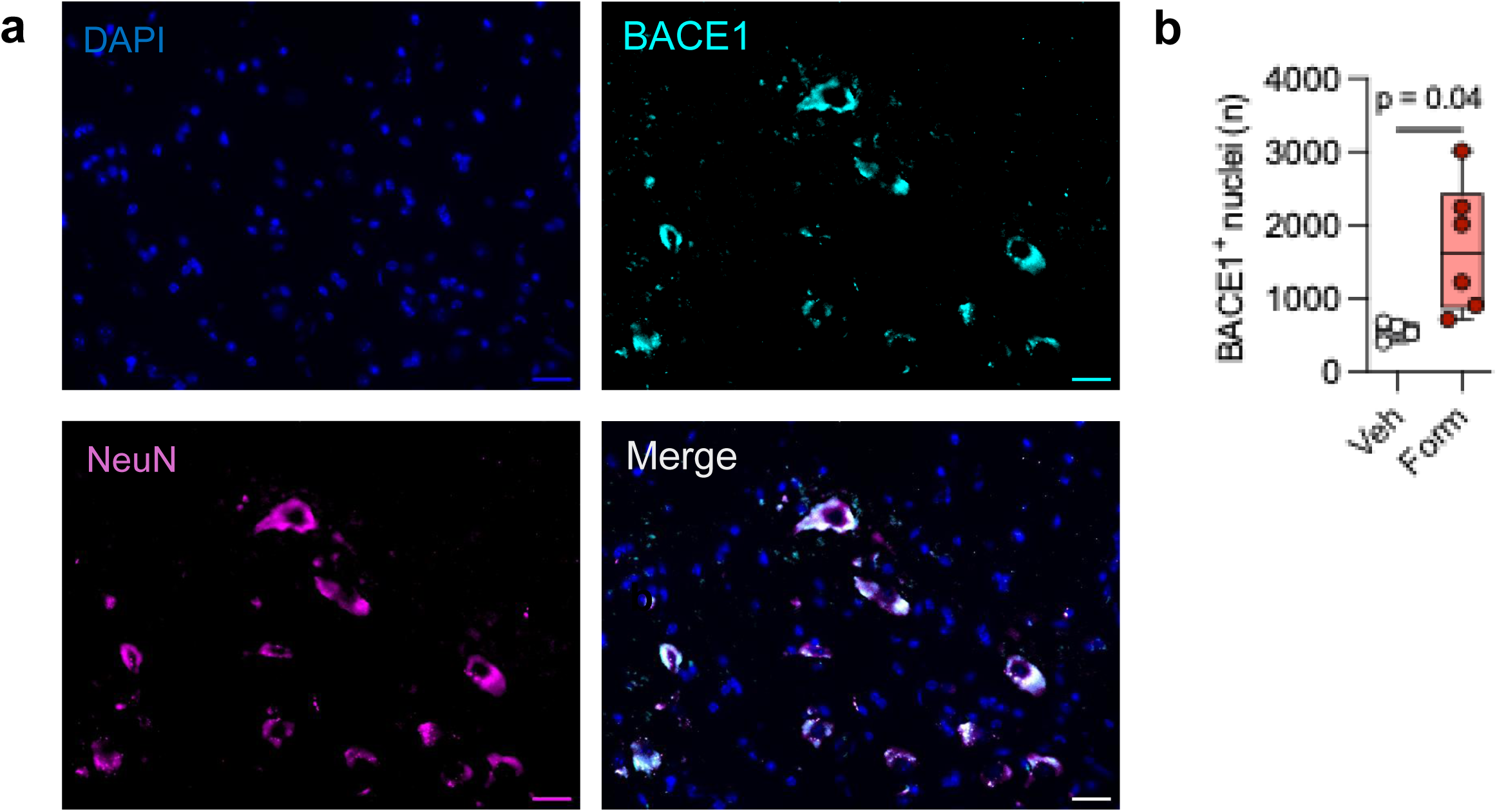

**Figure S10.**
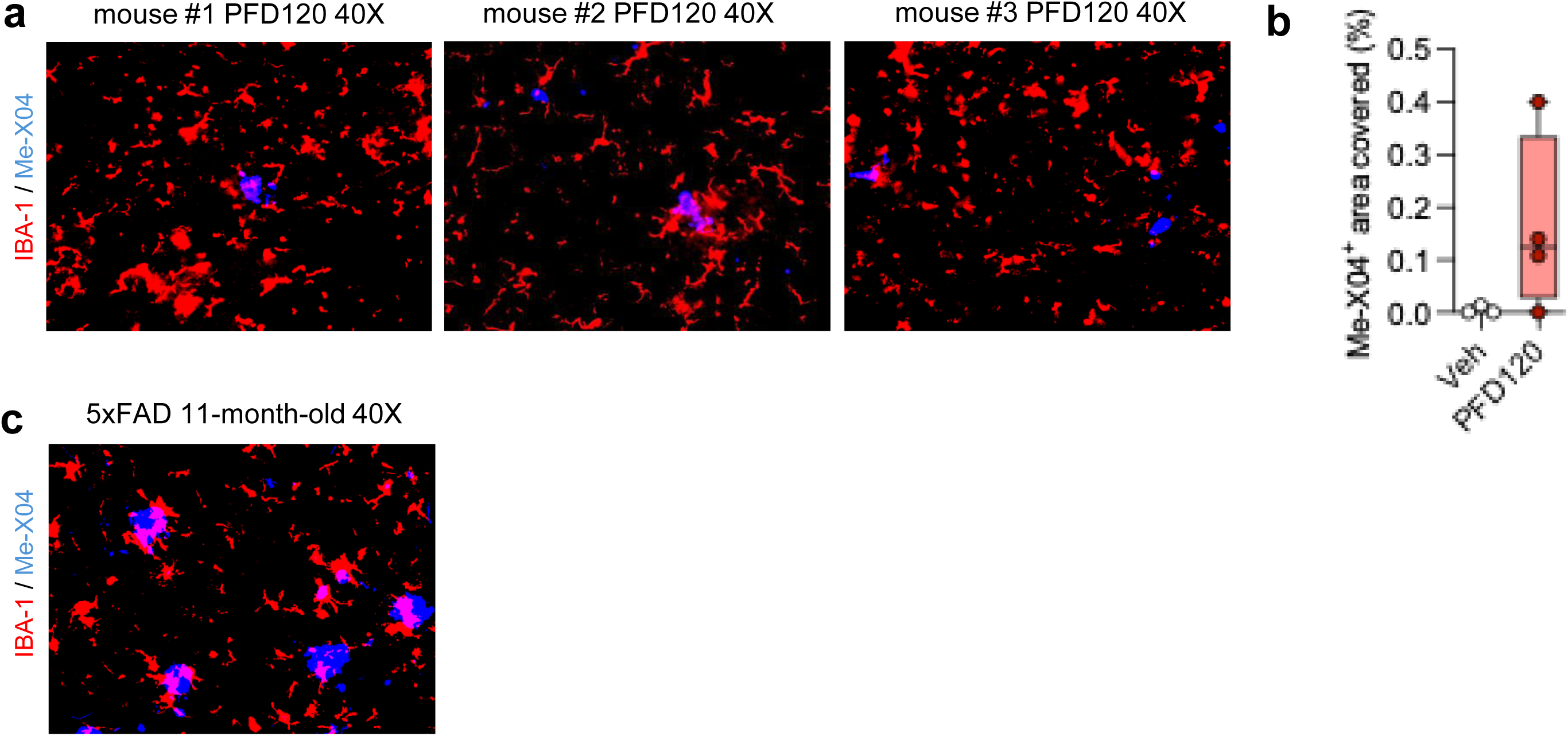

**Figure S11.**
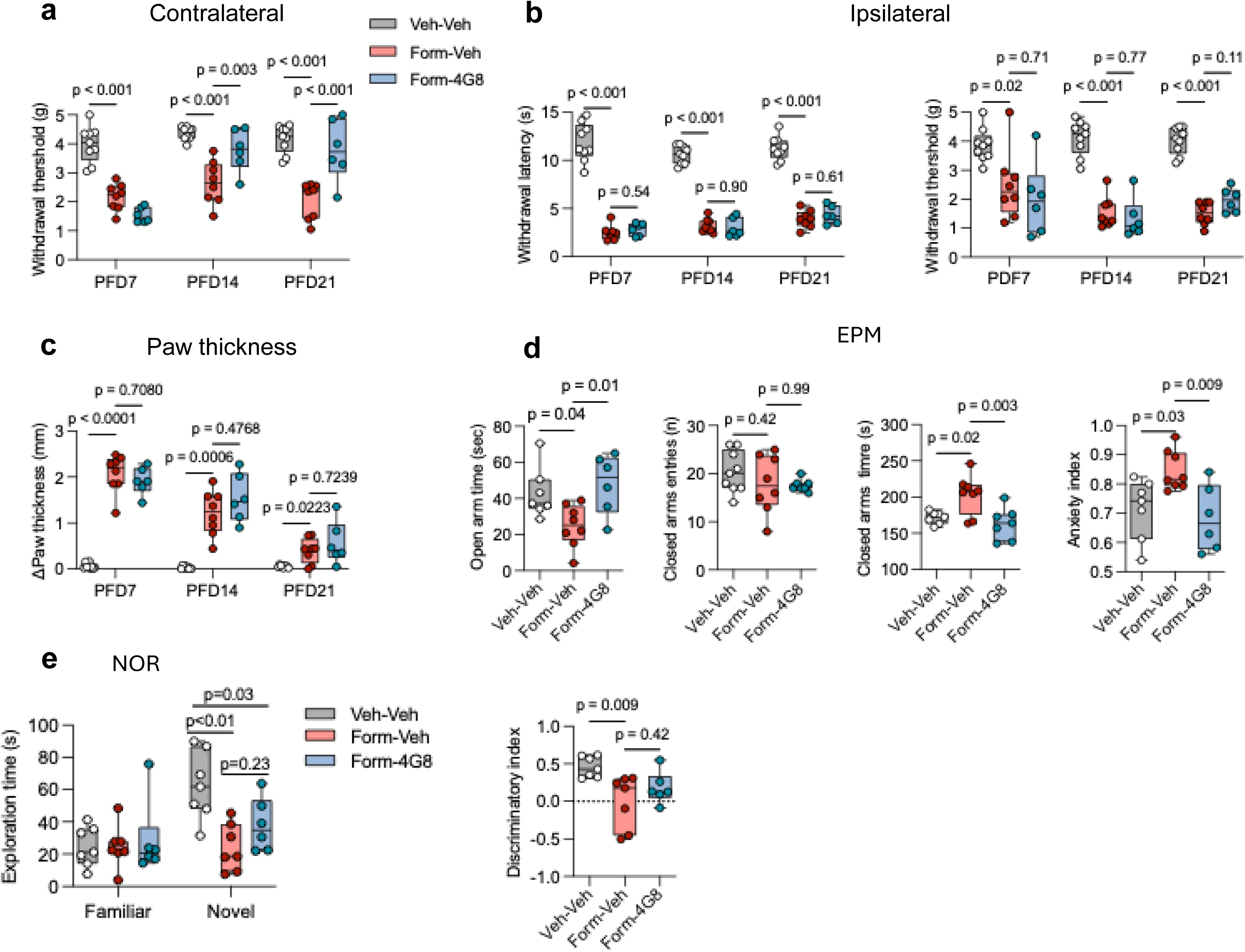

**Figure S12.**
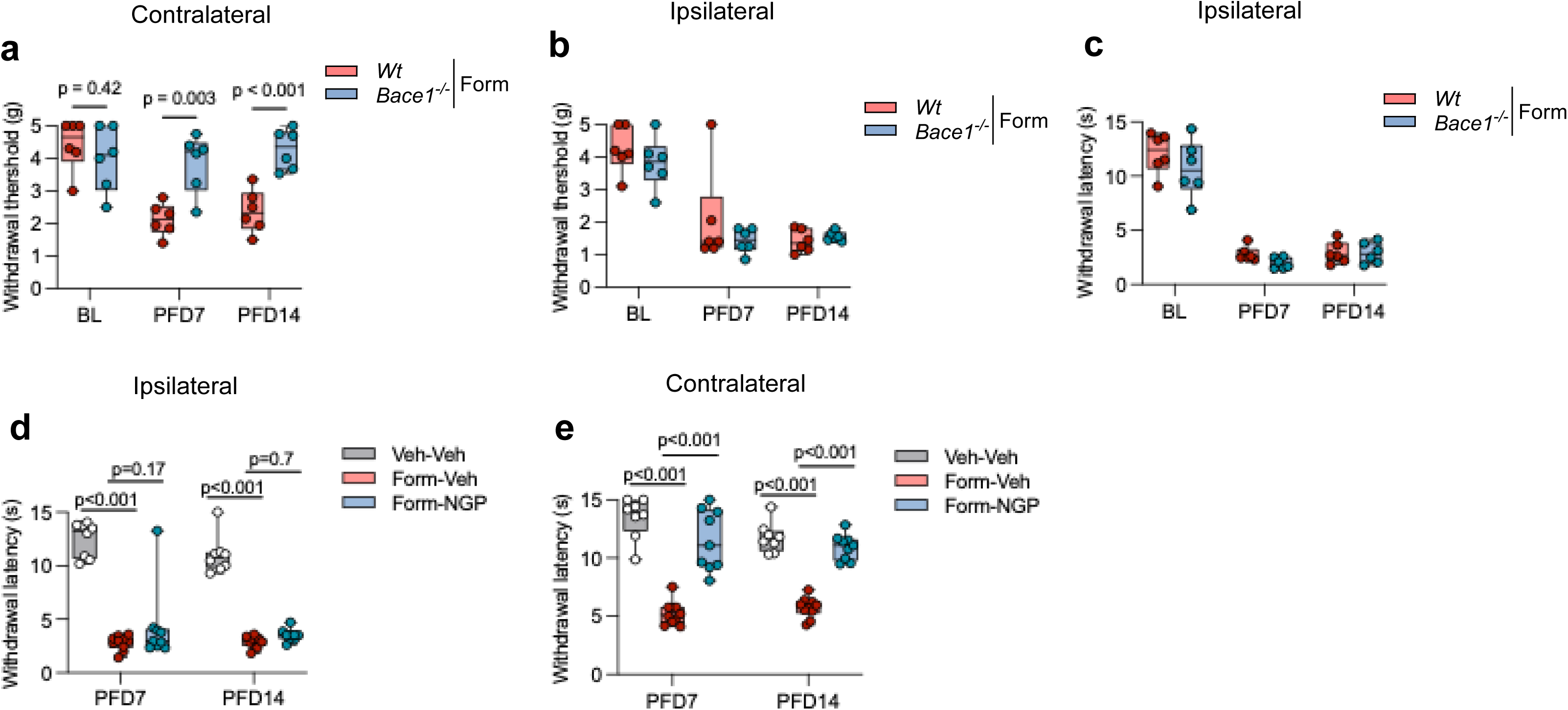

**Figure S13.**
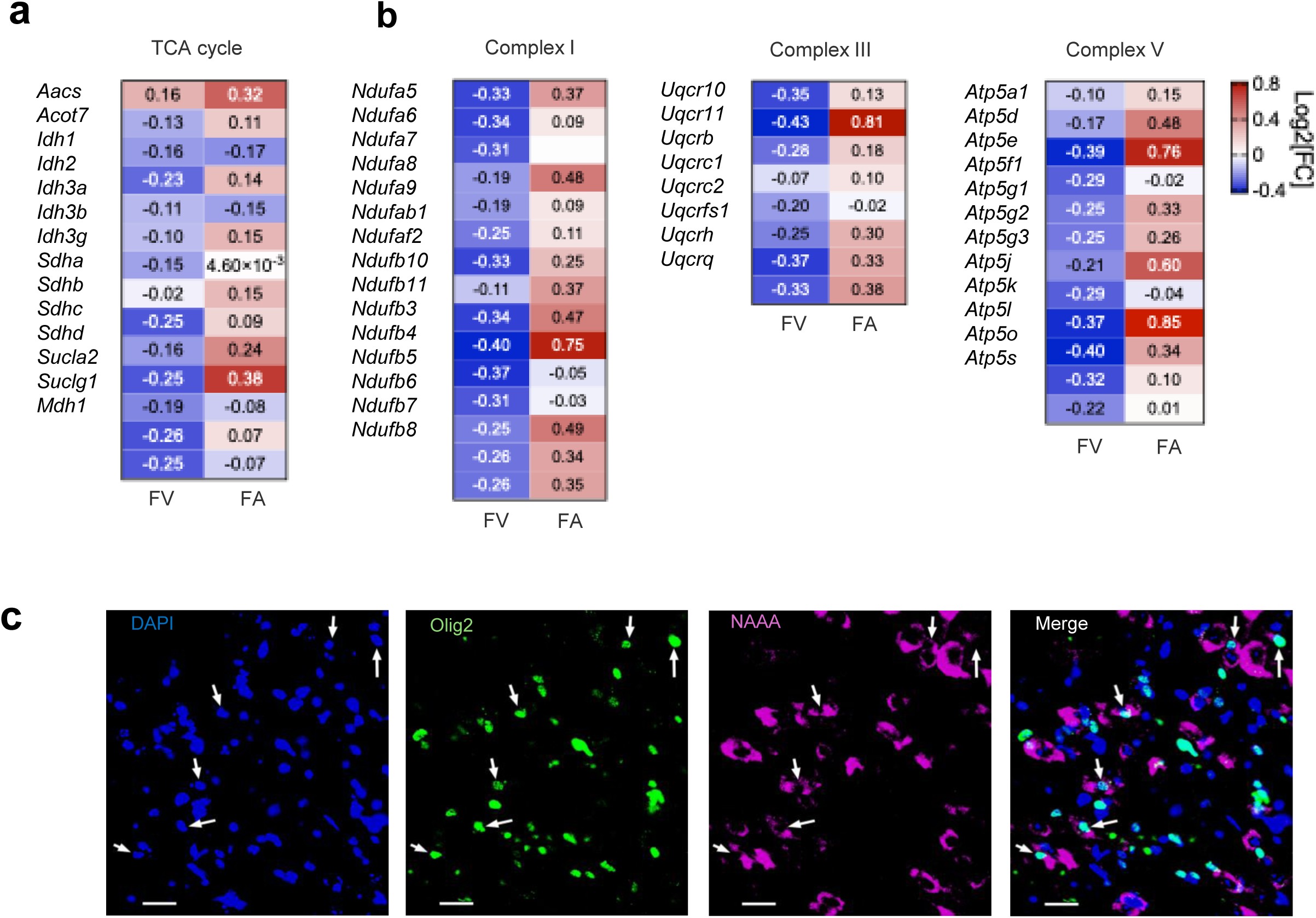

**Figure S14.**
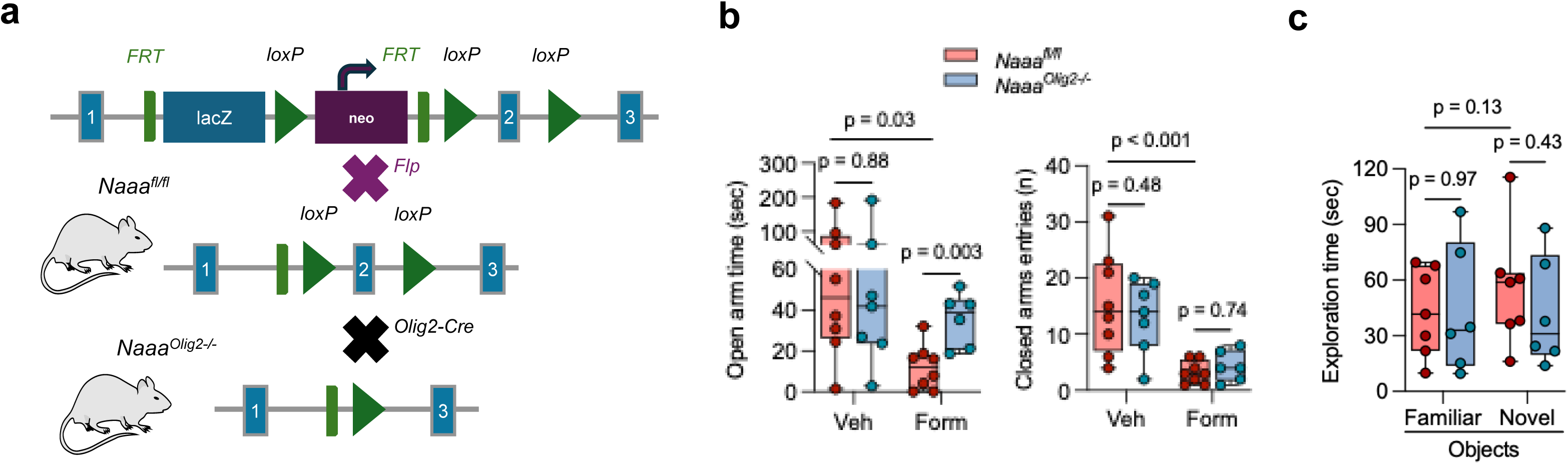

**Figure S15.**
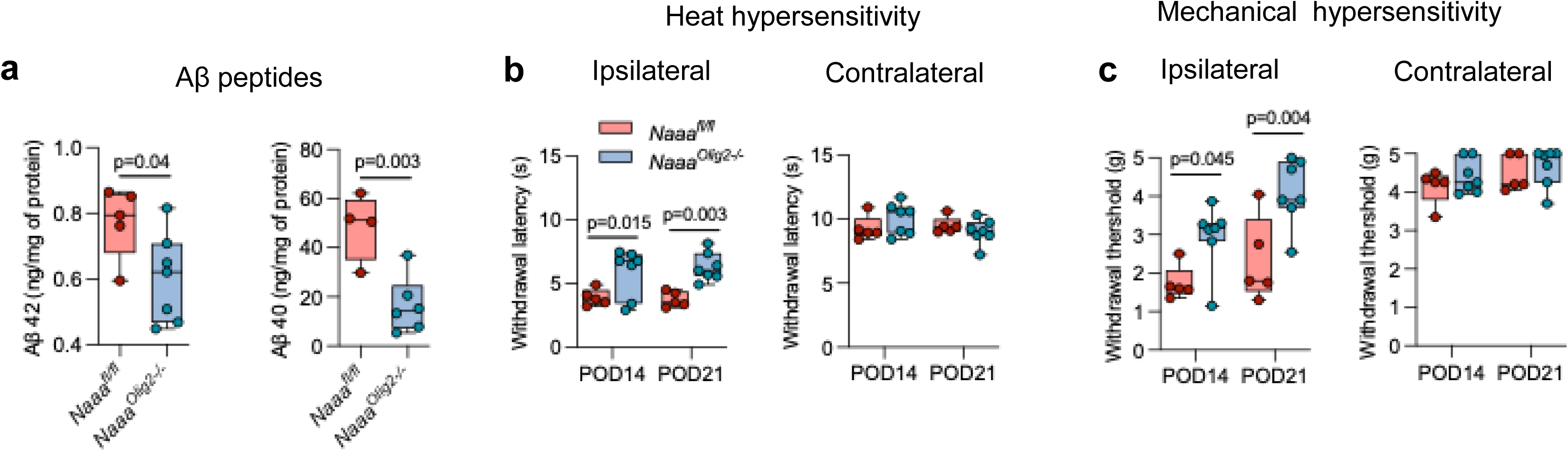

## References

1. P. Lavand’homme, Transition from acute to chronic pain after surgery. Pain 158 Suppl 1, S50–S54 (2017).

2. I. A. Harris, J. M. Young, H. Rae, B. B. Jalaludin, M. J. Solomon, Factors associated with back pain after physical injury: a survey of consecutive major trauma patients. Spine (Phila Pa 1976) 32, 1561–1565 (2007).

3. R. Kuner, H. Flor, Structural plasticity and reorganisation in chronic pain. Nat Rev Neurosci 18, 113 (2017).

4. M. N. Baliki, A. V. Apkarian, Nociception, Pain, Negative Moods, and Behavior Selection. Neuron 87, 474–491 (2015).

5. A. I. Basbaum, D. M. Bautista, G. Scherrer, D. Julius, Cellular and molecular mechanisms of pain. Cell 139, 267–284 (2009).

6. G. Sideris-Lampretsas, M. Malcangio, Microglial heterogeneity in chronic pain. Brain Behav Immun 96, 279–289 (2021).

7. R. R. Ji, C. R. Donnelly, M. Nedergaard, Astrocytes in chronic pain and itch. Nat Rev Neurosci 20, 667–685 (2019).

8. Y. Fotio et al., NAAA-regulated lipid signaling in monocytes controls the induction of hyperalgesic priming in mice. Nat Commun 15, 1705 (2024).

9. Y. Fotio et al., NAAA-regulated lipid signaling governs the transition from acute to chronic pain. Sci Adv 7, eabi8834 (2021).

10. A. M. Tagne et al., Metabolic reprogramming in the spinal cord drives the transition to pain chronicity. bioRxiv, (2025).

11. D. Piomelli et al., N-Acylethanolamine Acid Amidase (NAAA): Structure, Function, and Inhibition. J Med Chem 63, 7475–7490 (2020).

12. J. Fu et al., Oleylethanolamide regulates feeding and body weight through activation of the nuclear receptor PPAR-alpha. Nature 425, 90–93 (2003).

13. J. Lo Verme et al., The nuclear receptor peroxisome proliferator-activated receptor-alpha mediates the anti-inflammatory actions of palmitoylethanolamide. Mol Pharmacol 67, 15–19 (2005).

14. C. R. Bartman, T. TeSlaa, J. D. Rabinowitz, Quantitative flux analysis in mammals. Nat Metab 3, 896–908 (2021).

15. C. A. Porro et al., Effects of ketamine anesthesia on central nociceptive processing in the rat: a 2-deoxyglucose study. Neuroscience 125, 485–494 (2004).

16. B. Valerio-Gomes, D. M. Guimaraes, D. Szczupak, R. Lent, The Absolute Number of Oligodendrocytes in the Adult Mouse Brain. Front Neuroanat 12, 90 (2018).

17. F. Akbik, W. B. Cafferty, S. M. Strittmatter, Myelin associated inhibitors: a link between injury-induced and experience-dependent plasticity. Exp Neurol 235, 43–52 (2012).

18. Y. Poitelon, A. M. Kopec, S. Belin, Myelin Fat Facts: An Overview of Lipids and Fatty Acid Metabolism. Cells 9, (2020).

19. E. T. Walters, R. J. Crook, G. G. Neely, T. J. Price, E. S. J. Smith, Persistent nociceptor hyperactivity as a painful evolutionary adaptation. Trends Neurosci 46, 211–227 (2023).

20. W. Y. Aung, S. Mar, T. L. Benzinger, Diffusion tensor MRI as a biomarker in axonal and myelin damage. Imaging Med 5, 427–440 (2013).

21. A. Frati et al., Diffuse Axonal Injury and Oxidative Stress: A Comprehensive Review. Int J Mol Sci 18, (2017).

22. K. Sato, K. I. Takayama, M. Hashimoto, S. Inoue, Transcriptional and Post-Transcriptional Regulations of Amyloid-beta Precursor Protein (APP) mRNA. Front Aging 2, 721579 (2021).

23. J. Koenigsknecht, G. Landreth, Microglial phagocytosis of fibrillar beta-amyloid through a beta1 integrin-dependent mechanism. J Neurosci 24, 9838–9846 (2004).

24. C. Depp et al., Myelin dysfunction drives amyloid-beta deposition in models of Alzheimer’s disease. Nature 618, 349–357 (2023).

25. P. Yuan et al., TREM2 Haplodeficiency in Mice and Humans Impairs the Microglia Barrier Function Leading to Decreased Amyloid Compaction and Severe Axonal Dystrophy. Neuron 90, 724–739 (2016).

26. C. Balducci et al., Synthetic amyloid-beta oligomers impair long-term memory independently of cellular prion protein. Proc Natl Acad Sci U S A 107, 2295–2300 (2010).

27. N. Shenker et al., A review of contralateral responses to a unilateral inflammatory lesion. Rheumatology (Oxford) 42, 1279–1286 (2003).

28. L. Arendt-Nielsen, Central sensitization in humans: assessment and pharmacology. Handb Exp Pharmacol 227, 79–102 (2015).

29. M. C. Bushnell, M. Ceko, L. A. Low, Cognitive and emotional control of pain and its disruption in chronic pain. Nat Rev Neurosci 14, 502–511 (2013).

30. P. C. May et al., Robust central reduction of amyloid-beta in humans with an orally available, non-peptidic beta-secretase inhibitor. J Neurosci 31, 16507–16516 (2011).

31. J. D. Scott et al., Discovery of the 3-Imino-1,2,4-thiadiazinane 1,1-Dioxide Derivative Verubecestat (MK-8931)-A beta-Site Amyloid Precursor Protein Cleaving Enzyme 1 Inhibitor for the Treatment of Alzheimer’s Disease. J Med Chem 59, 10435–10450 (2016).

32. K. D. Rynearson et al., Preclinical validation of a potent gamma-secretase modulator for Alzheimer’s disease prevention. J Exp Med 218, (2021).

33. Y. Fotio, O. Sasso, R. Ciccocioppo, D. Piomelli, Antinociceptive Profile of ARN19702, (2-Ethylsulfonylphenyl)-[(2S)-4-(6-fluoro-1,3-benzothiazol-2-yl)-2-methylpiperazin-1-yl]methanone, a Novel Orally Active N-Acylethanolamine Acid Amidase Inhibitor, in Animal Models. J Pharmacol Exp Ther 378, 70–76 (2021).

34. T. J. Coderre, A. Laferriere, The emergence of animal models of chronic pain and logistical and methodological issues concerning their use. J Neural Transm (Vienna) 127, 393–406 (2020).

35. S. Gritsch et al., Oligodendrocyte ablation triggers central pain independently of innate or adaptive immune responses in mice. Nat Commun 5, 5472 (2014).

36. M. Simons, E. M. Gibson, K. A. Nave, Oligodendrocytes: Myelination, Plasticity, and Axonal Support. Cold Spring Harb Perspect Biol 16, (2024).

37. A. Latremoliere, C. J. Woolf, Central sensitization: a generator of pain hypersensitivity by central neural plasticity. J Pain 10, 895–926 (2009).

38. C. Fruhbeis et al., Oligodendrocytes support axonal transport and maintenance via exosome secretion. PLoS Biol 18, e3000621 (2020).

39. E. M. Kramer-Albers, H. B. Werner, Mechanisms of axonal support by oligodendrocyte-derived extracellular vesicles. Nat Rev Neurosci 24, 474–486 (2023).

40. E. Asadollahi et al., Oligodendroglial fatty acid metabolism as a central nervous system energy reserve. Nat Neurosci 27, 1934–1944 (2024).

41. C. W. Mount, M. Monje, Wrapped to Adapt: Experience-Dependent Myelination. Neuron 95, 743–756 (2017).

42. M. Kenigsbuch et al., A shared disease-associated oligodendrocyte signature among multiple CNS pathologies. Nat Neurosci 25, 876–886 (2022).

43. W. Fan, R. Evans, PPARs and ERRs: molecular mediators of mitochondrial metabolism. Curr Opin Cell Biol 33, 49–54 (2015).

44. C. Titus, M. T. Hoque, R. Bendayan, PPAR agonists for the treatment of neuroinflammatory diseases. Trends Pharmacol Sci 45, 9–23 (2024).

45. S. Wojtowicz, A. K. Strosznajder, M. Jezyna, J. B. Strosznajder, The Novel Role of PPAR Alpha in the Brain: Promising Target in Therapy of Alzheimer’s Disease and Other Neurodegenerative Disorders. Neurochem Res 45, 972–988 (2020).

46. N. Bornier et al., Chronic pain is a risk factor for incident Alzheimer’s disease: a nationwide propensity-matched cohort using administrative data. Front Aging Neurosci 15, 1193108 (2023).

47. J. Tian et al., Association between chronic pain and risk of incident dementia: findings from a prospective cohort. BMC Med 21, 169 (2023).

48. T. R. Bell et al., Multisite chronic pain: association with cognitive decline and post-mortem Alzheimer’s biomarkers. Brain Commun 7, fcaf208 (2025).

49. P. Branco et al., Structural brain connectivity predicts early acute pain after mild traumatic brain injury. Pain 164, 1312–1320 (2023).

