## Supplementary material for "Metabolic reallocation in spinal cord oligodendrocytes drives chronic pain via neuronal β-amyloid production": methods and material

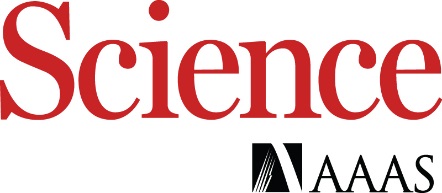

**Metabolic reallocation in spinal cord oligodendrocytes**

**drives chronic pain via neuronal β-amyloid production**

Yannick Fotio^1#^, Saeed Al Masri^1,4^, Zechuan Shi^2^, Johnny Le^3^, Sudeshna Das^2^, Varvara I. Rubtsova^3^,

Alex Mabou Tagne^1^, Cholsoon Jang^3^, Vivek Swarup^2^, and Daniele Piomelli^1,3,4, &^

^1^Department of Anatomy and Neurobiology, University of California Irvine, Irvine, CA, USA.

^2^Department of Neurobiology and Behavior, University of California, Irvine, Irvine, CA, USA

^3^Department of Biological Chemistry, University of California Irvine, Irvine, CA, USA.

^4^Department of Pharmaceutical Sciences, University of California Irvine, Irvine, CA, USA

^#^ Current address: Department of Biological Sciences, Center for Advances Pain Studies, The University of Texas at Dallas, Dallas, TX, USA

**This PDF file includes:**

Materials and Methods

Supplementary Text

Figs. S1 to S15

**Materials and Methods**

**Chemicals**

LY2811376, MK-8931, and NGP555 were purchased from Selleckchem (San Diego, CA) and dissolved in a vehicle of dimethyl sulfoxide (DMSO)/polyethylene glycol (PEG)-400/Tween 80/distilled water (5/40/5/50, vol). [U-^13^C]-glucose was from Cambridge Isotope Laboratories (Tewksbury, MA) and was dissolved in saline. ARN19702 was synthesized according to established protocols (*1*) and dissolved in PEG-400/Tween 80/distilled water (15/15/70, vol). The monoclonal anti-amyloid β antibody 4G8 (cat # 800710) was purchased from BioLegend (San Diego, CA). Methoxy-X04 was from Tocris (cat. # 4920) and dissolved in a vehicle of dimethyl sulfoxide (DMSO)/polyethylene glycol (PEG)-400/Tween 80/distilled water (5/40/5/50, vol). All other solvents and reagents were of the highest grade available.

**Study approval**

All experimental procedures complied with the ethical regulations for the care and use of laboratory animals promulgated by the National Institutes of Health (NIH) and the International Association for the Study of Pain (IASP). Formal approval was obtained from the Animal Care and Use Committee of the University of California, Irvine (AUP-23-082).

**Animals**

We used C57Bl/6 mice (~25 g, 10-12 weeks old; Jackson Laboratory, CA) unless otherwise indicated. Global *Naaa*-null (*Naaa^−/−^*) mice were purchased from MRC Harwell (Didcot, UK) via the EMMA-European Mouse Mutant Archive. A detailed description of their generation and phenotypic characterization may be found in (*2, 3*). Oligodendrocyte-specific *Naaa*-null (*Naaa^Olig2−/−^*) mice were obtained as follows: NAAA-floxed mice (*Naaa^fl/fl^*) mice were generated by mating *Naaa^-/-^* mice with flippase (Flp) FLPo-10 mice (Jackson Lab, #011065) and cross-bred with *Olig2^Cre^* (Jackson Lab, # 025567) to yield *Naaa^Olig2−/−^* mice, which were viable and fertile. Heterozygous *Bace1*-null *(Bace1^−/+^)* mice were purchased from Jackson Laboratories (#004714). Their generation and phenotypic profile were previously reported (*4*). 5×FAD hemizygous (*B6.Cg-Tg(APPSwFlLon,PSEN1*M146L*L286V)6799Vas/Mmjax*) mice and their wild-type littermates were a gift of Dr. Kim Green (University of California, Irvine). All mice were maintained on a C57Bl/6 genetic background. Genotyping was performed at TransnetYX (Duarte, CA) on earmark-derived tissue samples. Mice were group-housed under a 12-hour light/dark cycle at controlled temperature (20 ± 2°C) and humidity (55-60%) with food and water available *ad libitum*. They were randomly assigned to treatment groups and, before the start of the experiments, were handled for 3 consecutive days (~3 minutes per animal/day). Behavioral testing was conducted during the light phase of the light/dark cycle. Efforts were made to minimize the number of animals used and their discomfort.

**Drug administration**

Drug solutions and vehicles were prepared immediately before use and administered via either intraperitoneal, oral, or intrathecal administration. Formalin (1% in sterile saline, 20 μL; Sigma-Aldrich, St. Louis, MO) was administered by subcutaneous injection into the mouse right hind paw.

**Metabolic flux**

Four days after formalin injection, we administered [U-^13^C]-glucose (1 g/kg in saline) by oral gavage. Twenty minutes later, we collected ipsilateral L4-L6 spinal cord segments and snap froze them in liquid nitrogen. Tissue samples (∼20 mg each) were weighed on dry ice and homogenized in a mixture of cold methanol/acetonitrile/water (40/40/20, vol; 1-1.2 ml). The homogenates were vortexed and centrifuged at 16,000 × *g* for 10 minutes at 4°C.

**Myelin fractionation**

Myelin was partially purified using sucrose density gradient centrifugation, as described (*5*), with small modifications. Lumbar L4-L6 spinal cord fragments (~20 mg each) were transferred into 2-ml Precellys soft-tissue tubes (Bertin Instruments, France) and suspended in a sucrose solution (0.3 M) containing phosphatase (Sigma Millipore, Cat #P5726) and protease (Thermo Fisher, Cat #78429) inhibitors. Samples were homogenized at 4°C using a Precellys homogenizer at 6000 rpm for 15 seconds. The homogenates were layered over sucrose (0.83 M) in 2-ml centrifuge tubes (Beckman Coulter, Brea, CA) and ultracentrifuged at 75,000 × *g* for 35 minutes at 4°C. The crude myelin band formed at the 0.3 M/0.83 M sucrose interface was collected. Total protein concentration was determined using the bicinchoninic acid (BCA) assay (Bio-Rad Laboratories, Hercules, CA).

**Immunofluorescence**

Animals were anesthetized with isoflurane and perfused transcardially with ice-cold phosphate-buffered saline (PBS), followed by 4% paraformaldehyde (PFA) in PBS (pH 7.4). Spinal cords were extruded, post-fixed for 4 hours in 4% PFA, and cryoprotected overnight in 30% sucrose at 4°C. After being embedded in optimal cutting temperature medium (Tissue-Tek^®^, Sakura Finetek, Torrance, CA), they were flash-frozen in cold isopentane. Three series of transversal sections (thickness: 10 μm) were prepared using a cryostat. After rinsing with 0.1 M PBS, the sections were blocked and permeabilized with 0.3% Triton-X in 0.1 M PBS containing 3% normal horse serum for 60 minutes at room temperature. Single and double immunostaining experiments were performed by incubating tissues overnight at 4°C with primary antibodies [anti-NAAA (1:100; Invitrogen, Carlsbad, CA, Cat. # PA5-69357); anti-IBA1 (1:1000; Abcam, Waltham, MA, Cat. # ab5076); anti-Olig2 (1:200; Millipore, Hayward, CA, #MABN50); anti-APP (1:1000; Millipore, Hayward, CA, #EMD171610); anti-Aβ (clone 4G8, 1:1000, Biolegend, San Diego, CA, Cat. # 800712)], followed by incubation for one hour at room temperature with appropriate secondary Alexa Fluor antibodies (1:1000; Invitrogen). Slides were mounted with fluorescence-safe mounting medium containing 4′,6-diamidino-2-phenylindole (DAPI) for nuclear staining. Images were obtained at 10x or 40x magnification using a BZ-X710 fluorescence microscope coupled with BZ-X Analyzer software (Keyence, Itasca, IL). Quantification of fluorescence intensity was performed using the NIH Fuji Is Just ImageJ (FIJI) software.

**In vivo amyloid-β staining**

On post-formalin day 120, C57BL/6J mice were injected intraperitoneally with methoxy-XO4 (10 mg/kg, body weight). After 24 hours, they were transcardially perfused with ice-cold PBS followed by 4% PFA. Spinal cord was extruded and processed for immunofluorescence as described above, with the exception that slides were mounted using a DAPI-free, fluorescence-safe mounting medium. Naïve age-matched 5×FAD mice (11 months old) were used as positive controls.

**Aβ measurement by enzyme-linked immunosorbent assay**

L4–L6 spinal cord fragments (~20 mg each) were placed into 2-mL Precellys soft tissue homogenization tubes (Bertin Instruments) and suspended in T-PER buffer (Thermo Fisher, #78510) supplemented with 1× phosphatase inhibitor (Sigma Millipore, #P5726) and protease inhibitor cocktails (Thermo Fisher, #78429). Samples were homogenized at 4°C using a Precellys homogenizer at 6000 rpm for two 15-second cycles, with a 20-second pause between cycles. The homogenates were transferred to 2-mL ultracentrifuge tubes (Beckman Coulter) and subjected to ultracentrifugation at 119,045 × *g* for 60 minutes at 4°C. The resulting supernatants were collected as the soluble fraction. The pellets were resuspended into a formic acid : distilled water solution (70:30, v/v), homogenized, and ultracentrifuged under the same conditions. The resulting supernatants were collected as the insoluble fraction. Amyloid-β (Aβ40 and Aβ42) concentrations were quantified in both soluble and insoluble fractions using ELISA kits (Thermo Fisher Scientific, #KHB3481 and #KHB3441) following manufacturer’s instructions.

**Myelin fractionation**

Myelin was extracted using sucrose density gradient centrifugation, as described (*5*), with small modifications. Lumbar L4-L6 spinal cord fragments (~20 mg each) were transferred into 2-ml Precellys soft-tissue tubes (Bertin Instruments, France) and suspended in a sucrose solution (0.3 M) containing phosphatase (Sigma Millipore, Cat. #P5726) and protease (Thermo Fisher, Cat. #78429) inhibitors. Samples were homogenized at 4°C using a Precellys homogenizer at 6000 rpm for 15 seconds. The homogenates were layered over sucrose (0.83 M) in 2-ml centrifuge tubes (Beckman Coulter) and ultracentrifuged at 75,000 × *g* for 35 minutes at 4°C. The crude myelin band formed at the 0.3 M/0.83 M sucrose interface was collected. Total protein concentration was determined using the bicinchoninic acid (BCA) assay (Bio-Rad Laboratories, Hercules, CA).

**Metabolic flux studies**

*Sample preparation*

We injected formalin (1%, 20 μL) or its vehicle (saline) in the hind-paw of male mice (n = 4 per group) and, after four days, administered [U-^13^C]-glucose (1 g/kg in saline) via the oral route. Twenty minutes later, we collected ipsilateral L4-L6 spinal cord segments and snap-froze them in liquid nitrogen. Tissue samples (∼20 mg each) were weighed on dry ice and homogenized in a mixture of cold methanol/acetonitrile/water (40/40/20, vol; 1-1.2 ml). The homogenates were vortexed and centrifuged at 16,000 × *g* for 10 minutes at 4°C. The supernatants (3 μL) were subjected to liquid chromatography-mass spectrometry (LC/MS) analysis, as detailed below.

*Liquid chromatography-mass spectrometry (LC/MS)*

The same LC-MS method was applied to both myelin lipidomic and metabolic flux analyses. Samples (3 μL) were analyzed using a quadrupole orbitrap mass spectrometer (Q Exactive Plus, Thermo Fisher Scientific, San Jose, CA) operating in negative or positive ion modes, coupled to a Vanquish Ultra High-Performance LC system (UHPLC, Thermo Fisher Scientific) with electrospray ionization and used to scan *m/z* 70 to 1000. Scanning frequence was 2 Hz and resolution was 140,000. LC separation was achieved on a XBridge BEH Amide column (2.1 mm x 150 mm, 2.5 mm particle size, 130 Å pore size, Waters) using a gradient consisting of solvent A (20 mM ammonium acetate, 20 mM ammonium hydroxide in 95:5 water: acetonitrile, pH 9.45) and solvent B (acetonitrile). The gradient was: 0 min, 85% B; 2 min, 85% B; 3 min, 80% B; 5 min, 80% B; 6 min, 75% B; 7 min, 75% B; 8 min, 70% B; 9 min, 70% B; 10 min, 50% B; 12 min, 50% B; 13 min, 25% B; 16 min, 25% B; 18 min, 0% B; 23 min, 0% B; 24 min, 85% B; 30 min, 85% B. Flow rate was 150 μL/min. Autosampler temperature was 5°C.

*Data analysis*

Data were converted from *.raw* files to *.mzXML* and then loaded into EI-MAVEN open-source software (Version 0.12.0, Elucidata). Using an annotated MS library of retention times and accurate *m/z* values generated in house, we selected metabolite peaks of interest on EI-MAVEN. Peak shape and quality were evaluated and Area Top of the metabolites of interest was recorded. For isotope tracing experiment, natural isotope correction was performed with AccuCor2 R code (https://github.com/wangyujue23/AccuCor2) and IsoCorrectoR. Microsoft Excel and R were used for data analysis.

**Proteomics**

*Sample Preparation*

Myelin extracts and L4–L6 spinal cord segments were snap-frozen and stored at -80°C. Protein sample preparation was carried out using an SP3 (Single-Pot, Solid-Phase-enhanced Sample Preparation) approach. Briefly, proteins were extracted in 100 mM ammonium bicarbonate (ABC) containing 1% SDS and homogenized using a Beadbeater (BioSpec Products, Bartlesville, OK) for 4 cycles of 30 seconds separated by ice cooling. Total protein concentration was measured using the BCA assay. Following dilution to a consistent protein concentration of 1 ug/uL with 1% SDS in 100 mM ABC, a fraction of each sample was pooled and split into three technical replicates. 50 µg of protein in 50 µL of lysis buffer per sample was reduced with 10 mM dithiothreitol (DTT) for 15 minutes at 65°C, then alkylated with 20 mM iodoacetamide (IAA) at room temperature in the dark. Pre-washed SP3 paramagnetic beads (1:1 hydrophilic/hydrophobic mix; Cytiva, Pensacola, FL) were added at a bead-to-protein ratio of 20:1 (w/w). Acetonitrile was then added to reach a final concentration of 50% (v/v), and the samples were incubated for 20 minutes at room temperature. The tubes were placed on a magnetic rack for 2 minutes, and the supernatants were carefully removed. The beads were washed by performing two cycles of resuspending in 80% ethanol (1 ml) followed by magnetic separation and aspiration, then once in acetonitrile (1 ml). The bead-bound proteins were resuspended in 75 µL of 100 mM ABC. Proteolytic digestion was performed overnight at 37^o^C with continuous shaking (1,000 rpm, Thermomixer), using sequencing-grade trypsin (Promega, Madison, WI) at a 1:20 (w/w) enzyme-to-protein ratio. Digestion was quenched by adding formic acid to a final 1% (v/v) concentration, and the beads were magnetized. The supernatant was then transferred and diluted to a final concentration of 0.4 µg/µL (starting protein) with 0.1% formic acid in an injection vial for LC–MS/MS analysis.

*LC/MS-MS*

LC/MS-MS analyses were performed using a Thermo Vanquish UHPLC system coupled to a Q Exactive Plus mass spectrometer (Thermo Fisher Scientific) maintained using manufacturer protocols and checked for system suitability using Pierce HeLa digest standard. Each sample (20 µg in 50 µL) was loaded onto a Waters CSH C18 column (150 mm × 1 mm, 1.7 µm particle size) maintained at 55^o^C. Mobile phase A was water containing 0.1% formic acid, and mobile phase B was acetonitrile containing 0.1% formic acid. A 70-minute linear gradient was used at a flow rate of 70 µL/min from 0 to 50 minutes, increasing from 0% B at 0 minutes to 35% B at 50 minutes. The gradient then increased to 100% B at 60 minutes, followed by a re-equilibration phase at 0% B from 60 to 70 minutes at a flow rate of 140 µL/min. Top-15 data-dependent acquisition (DDA) was performed on the Q Exactive Plus in positive electrospray ionization mode with a narrow-bore HESI emitter set to 3.5 kV spray voltage, an S-lens setting of 50, a sheath gas flow of 50 units, a sweep gas flow of 1 unit, a capillary temperature of 300C, and an auxiliary heater temperature of 50C. MS1 scans were acquired at 70,000 resolution over an m/z range of 350–1650, with a normalized AGC target of 5e6 and a maximum injection time of 50 ms. MS2 scans were acquired at 17,000 resolution with 1.6 m/z-wide isolation windows, a fixed first mass of 110 m/z, a normalized HCD collision energy of 28%, an AGC target of 5e4, and a maximum injection time of 50 ms. The intensity threshold was set at 1e4, the charge inclusion range was 2 to 5, and dynamic exclusion was set to 30 seconds, with isotope exclusion enabled and peptide match preferred. All scans were acquired in centroid mode.

*Data analysis*

Relative protein quantification was carried out using MSFragger LFQ-MBR with the UniProt mouse reference proteome and potential contaminants, using default settings. Differential expression analysis (using the limma R package) and pathway enrichment analysis (using Kolmogorov–Smirnov tests) were performed in custom R scripts with Gene Ontology, Reactome, and WikiPathways mouse set databases. Significance was determined using the Benjamini–Hochberg FDR method for multiple testing correction.

**snRNA-seq**

Approximately 50 mg of freshly frozen L4-L6 spinal cord ipsilateral to the formalin injection site was prepared for snRNA-seq. Each tissue sample was harvested on post-formalin day 4 (PFD4) from three C57BL/6J mice, pooled to generate a single biological replicate, and homogenized in EZ lysis buffer (Sigma-Aldrich, Cat #NUC101–1KT) on ice for 10 minutes. The homogenates were filtered through a 70 μm mesh and transferred to a clean tube. After centrifugation at 500 x *g* for 5 minutes at 4°C, the pellet was resuspended in 1 ml of EZ lysis buffer, and the centrifugation step was repeated. The pellets were washed and incubated for 5 minutes in a buffer containing 1x PBS, 1% bovine serum albumin (BSA), and 0.2 U/μl RNase inhibitor to stabilize nuclei.

To remove myelin and other cellular debris we added a sucrose solution (1.8M) and the samples were centrifuged at 13,000 × *g* for 45 minutes at 4°C. Following this step, a debris removal solution (Miltenyi Biotec, Cat. #130-109-398) was added to the nuclei, and the mixture was centrifuged at 3,000 × *g* for 10 minutes at 4°C. The cleaned nuclei were diluted in buffer containing BSA and RNase inhibitor, then treated with the Nuclei Fixation Kit (Parse Biosciences Evercode™, Cat. # ECFN3300) for fixation and permeabilization. The nuclei were then cryopreserved in DMSO and stored until library preparation. Libraries were constructed using the EVERCODE™ WT V3 kit (Parse Biosciences, Cat. #ECWT3300). Quantification of DNA was performed with the Qubit dsDNA HS Assay Kit (Invitrogen, Cat #Q32851), and the average fragment size was determined with the D5000 HS Kit (Agilent, Cat #5067-5592 and #5067-5593). Sequencing was carried out on an Illumina Novaseq 6000 S4 system, using paired-end sequencing to achieve a target depth of 50,000 read pairs per nucleus.

*snRNA-seq data processing*

We processed snRNA-seq reads using the ParseBio Trailmarker pipeline (v1.4.0) to align fastq files to the GRCm39 reference transcriptome, downloaded from Ensembl, barcode and unique molecular identifiers (UMIs) extraction, and raw count gene matrix generation for each cell barcode. We then used Scrublet (v0.2.3) with default parameters to identify and exclude multiplets (barcodes associated with multiple nuclei). For initial QC analysis at database level, we excluded barcodes with more than 10,000 UMIs and fwer than 300 UMI and filtered out those with over 5% mitochondrial reads. At each sample level, we removed barcodes in the top 5% for UMI counts, and the top 10% of doublet scores. Clustering was performed using the Scanpy package, following these steps: (1) Normalizing gene expression by total UMI counts per cell and applying a logarithmic transformation using *sc.pp.normalize_total* and *sc.pp.log1p.* (2) Selecting highly variable genes with *sc.pp.highly_variable_genes*, using the “Seurat_v3” method, retaining 2,000 genes for downstream analyses. (3) Scaling the expression matrix for the selected genes to unit variance and zero mean with *sc.pp.scale*. (4) Performing principal component analysis (PCA) for dimensionality reduction using *sc.tl.pca*. (5) Constructing a cell-neighborhood graph based on the top 30 PCs using *sc.pp.neighbors* and visualizing it with UMAP via *sc.tl.umap*. (6) An initial Leiden clustering (resolution = 3) identified low-quality cell clusters that were removed based on canonical marker gene expression (*6*). To further validate cell type identities, we used *MapQuery* to project our dataset onto the reference mouse spinal cord snRNA-seq UMAP framework (*6*). After removing low-quality cells with predicted cell type score less than 0.9, we separated the dataset by major cell lineages for subclustering [excitatory neurons, inhibitory neurons, oligodendrocytes, astrocytes, oligodendrocyte precursor cells (OPC), motor neurons, ependymal cells, endothelial cells, cerebrospinal fluid-contacting neurons (CSF-cN), and vascular leptomeningeal cells (VLMC)]. This approach resulted in the final processed and clustered snRNA-seq dataset with 139,667 barcodes.

**Bulk RNA-seq**

Total RNA was extracted from ipsilateral lumbar spinal cord segments (L4–L6) using the RNeasy Mini Kit (Qiagen), and only samples with RNA integrity number (RIN) ≥8.5 were used for library preparation. cDNA synthesis, amplification, library construction, and sequencing were performed by Novogene (Beijing, China) on the Illumina NovaSeq platform using a 150 bp paired-end read configuration. Sequencing reads were aligned to the reference genome with STAR, and transcript abundance was quantified using HTSeq and Cufflinks. Differential gene expression analysis was performed with DESeq2 and edgeR, with genes showing adjusted P values < 0.05 considered significantly up- or downregulated.

**Transmission electron microscopy**

Specimens were fixed in 2% glutaraldehyde at 4^o^C for 12 hours, rinsed in neutral sodium cacodylate buffer, and post-fixed in 1.33% osmium oxide in cacodylate buffer for 2 hours at room temperature. The fixed samples were dehydrated through serial changes of 50 to 100% ethanol followed by propylene oxide. Tissues were infiltrated in a 1:1 mix of Epon 812 resin: propylene oxide for 60 minutes, then 100% Epon 812 for 60 minutes, then embedded in additional Epon 812 overnight. Post embedding, 1-mm thick sections were cut and counterstained with methylene blue /Azure II to confirm specimen orientation and quality. Ultra-thin (70-80 nm) sections were then cut onto grids, stained with 2% uranyl acetate for 10 minutes and then with Sato-modified lead stain for 3 minutes. Ultra-thin sections were imaged using an FEI Tecnai Spirit electron microscope. Images were analyzed using semi-automated software (MyelTracer) to determine the *g*-ratio, myelin thickness, and axonal diameter (*7*).

**Diffusion tensor imaging (DTI)**

Mice were anesthetized, flushed intracardially with ice-cold PBS and then with 4% PAF in PBS (pH 7.4). Spinal cords were removed within their osseous structures and post-fixed overnight in 4% PAF, followed by three 5-min washes with PBS the next day, and stored in PBS (pH 7.4) containing sodium azide (0.02%, weight) at 4^o^C. *Ex vivo* DTI studies were performed using a 9.4T Bruker Avance imager (Bruker Biospin, Billerica, USA). Data were acquired with the following parameters: 1.5 cm field of view, 0.5 mm slice thickness, and a 128x128 acquisition matrix and zero filled to 256x256. DTI parameters were as follows: repetition time (TR)/echo time (TE) =8000 ms/35.66 ms, 30 isolinear directions, B=3000 mT/m, and 5 B0 images acquired prior to weighted images. T2WI parameters were TR/TE=4000/10 ms with 10 equally spaced echoes. MRI scans were processed for eddy current correction as outlined above, and the scans were manually masked and extracted using ITK Snap (version 3.8.0). Masks were reviewed and adjusted by a blinded experimenter. FMRIB's Diffusion Toolbox was used to generate parametric DTI maps. The resultant maps were then analyzed in DSI studio (<http://dsi-studio.labsolver.org>). Distinct regions of interest throughout the spinal cord were drawn manually [see ref. (*8*) for accurate identification of spinal cord segments] for each animal without right/left distinction for white and gray matter. Regional average value of the diffusion indices (FA and MD) was automatically extracted for each spinal cord segment.

**Imaging mass spectrometry (IMS)**

Flash-frozen spinal cords were embedded in 2% carboxymethylcellulose and cryosectioned at 10 µm thickness onto Bruker MALDI IntelliSlides (Bruker Daltonics, Billerica, MA). The matrix 1,5,-diaminonaphthalene (DAN) was applied in multiple passes at 7.5 mg/mL in acetonitrile:water (70:30) for a final density of 120 µg/cm^2^ using an HTX M3+ matrix sprayer (HTX Technologies, Chapel Hill, NC). IMS data were acquired with a Shimadzu iMScope 9050 (AP-MALDI Q-TOF): MS^+^ mode, 100 laser shots at 5kHz, laser diameter setting 2, laser power 65.2%, pixel size 25 or 10 µm. IMDX data files were imported into Shimadzu ImageReveal software (Shimadzu Corp, Kyoto, Japan) for analysis.

**Pain models**

***Formalin test***

We adopted a variant of the formalin test developed to study the transition from acute to chronic pain (*2*). Formalin (1% v/v, 20 μl) or saline was administered by subcutaneous injection into the right hind paw. The mice were immediately transferred to a transparent observation chamber for behavioral analysis. Nocifensive responses, including the duration of licking or biting the injected paw and the frequency of paw flinches, were videorecorded over a 60-minute period and subsequently quantified by a blinded evaluator. On post-formalin days (PFD) 14 and 21, the following signs of pain chronification were assessed, as described below: (1) mechanical and thermal (heat) hypersensitivity in the ipsilateral (injured) and contralateral (non-injured) paws; (2) anxiety-like behavior in the elevated plus maze test; and (3) long-term memory in the novel-object recognition test.

***Chronic constriction injury (CCI)***

The CCI model was performed following an established protocol (*9*). Mice were anesthetized with isoflurane, and the right sciatic nerve was surgically exposed at the mid-thigh level using blunt dissection under aseptic conditions. The nerve was carefully cleared of surrounding connective tissue proximal to its trifurcation. Three chromic cat gut ligatures (4-0, Ethicon, Blue Ash, OH) were loosely tied around the nerve at 1-mm intervals to induce partial nerve constriction. The incision was closed with a single muscle suture and secured with skin clips. In sham-operated controls, the sciatic nerve was exposed but left untied.

**Behavioral tests**

***Thermal (heat) hypersensitivity***

Heat hypersensitivity was assessed using the Hargreaves plantar test apparatus (San Diego Instruments, San Diego, CA) as previously described (*2, 10, 11*). Following a 45-minute habituation period, the plantar surface of each hind paw was exposed to a radiant heat beam directed through the glass platform. A cut-off time of 15 seconds was implemented to prevent tissue damage. Each paw was tested three times, with a 2-minute interval between stimuli. The paw withdrawal latency (in seconds) was recorded for each trial and averaged to determine the response.

***Mechanical hypersensitivity***

Mechanical hypersensitivity was evaluated using a dynamic plantar aesthesiometer (Ugo Basile, Varese, Italy). The withdrawal threshold, defined as the force (in grams) required to elicit a paw withdrawal, was measured. A maximum force of 5 g was applied over 10 s as a cut-off. Baseline thresholds were determined through two consecutive measurements conducted two days prior to the administration of the priming agents.

***Elevated plus maze***

Each mouse was positioned on the central platform of the maze, oriented toward the open arm opposite the experimenter, and behavior was recorded using Debut video capture software (NCH Software, Canberra, Australia). A blind observer quantified the time spent in the open and closed arms, along with the number of entries into each. The open arms were illuminated at 150-170 lux, while the closed arms were maintained at 40-50 lux. The anxiety index was calculated as described (*2*).

***Novel object recognition***

The novel object recognition test was performed over a three-day period. On the first day, mice were acclimatized to an empty testing arena for 10 minutes. On the second day, two identical objects were introduced into the arena, and the mice were allowed to explore freely. On the third day, one of the original objects was replaced with a novel object differing in shape, color, and texture. Mice were given 10 minutes to explore the arena, during which the total time spent investigating each object (defined as nosing or sniffing within 2 cm of the object) was recorded by a blind observer. The discrimination index was calculated as previously reported (*2*).

**Supplemental figures**

**Figure S1: Effects of peripheral injury on cell type-specific gene transcription in the spinal cord.** (**a**) Distribution of nuclei across identified cell populations in ipsilateral L4-L6 spinal hemicords of male mice (n=4-7 per group) 4 days after intraplantar injection of formalin (red) or vehicle (cyan). (**b**) Heatmaps showing Log_2_ fold changes in expression of genes encoding components of mitochondrial complexes I, IV, and V. Red, upregulated; blue, downregulated. Abbreviations: Ast., astrocytes, CSFc, cerebrospinal fluid-contacting neurons; Exc., excitatory; Inh., inhibitory; M., motor; N., neurons; Olig., oligodendrocytes; OPC, oligodendrocyte precursor cells; VLMC, vascular leptomeningeal cells.

**Figure S2: Effects of peripheral injury on glucose transport and metabolism in the spinal cord.** (**a**) Log_2_ fold changes (formalin vs. vehicle) in expression of the glucose transporter type 1 (*Slc2a1*) in ipsilateral lumbar (L4-L6) spinal hemicords of male mice 4 days after injections. **(b, c)** [¹³C]-carbon incorporation into glycolysis metabolites (**b**) and TCA cycle metabolites (**c**) in ipsilateral L4-L6 spinal hemicords of male mice 4 days after injection of vehicle (gray symbols) or formalin (red symbols). **(d)** Pyruvate to citrate and pyruvate to malate ratios. Data in **(b, c)** are expressed as percentage of total [¹³C]-carbon incorporation.

*Statistical analysis*: Data were analyzed using unpaired *t*-tests (**a**). Box-and-whisker plots indicate the median and interquartile range. Individual data points represent mice (n=4-7 per group). *P* values reflect comparisons between formalin- and vehicle-treated groups.

**Figure S3: Effects of peripheral injury on spinal cord proteomics. (a)** Volcano plot illustrating differential protein expression in ipsilateral L4-L6 spinal hemicords of male mice 4 days after formalin or vehicle injection. Proteins significantly downregulated in formalin-treated mice are shown in blue, upregulated proteins are shown in red. (**b**) Quantification (Log_10_ ion count) of select proteins. Proteins involved in neuronal excitability and synaptic transmission: sodium channel protein type 2 subunit alpha (SCN2A), ATP-sensitive inward rectifier potassium channel 10 (KCNJ10), acetylcholine esterase (ACHE), glycine receptor subunit beta (GLRB). Proteins involved in synaptic vesicle trafficking and synaptogenesis: SV2B (synaptic vesicle glycoprotein 2B), VGAT (vesicular inhibitory amino acid transporter), SYN3 (synapsin 3), HTT (huntingtin), and SPARCL1 (secreted protein acidic and cysteine rich-like 1). Vehicle, open symbols; formalin, red symbols.

*Statistical analysis*: Data were analyzed using unpaired *t*-tests. Results are expressed as Log_10_ ion counts and displayed as box-and-whisker plots showing the median and interquartile range. Dots represent individual mice (n = 7 per group). *P* values versus vehicle are shown.

**Figure S4: Effects of peripheral injury on cell type-specific gene transcription in the spinal cord. (a)** Log_2_ fold changes (formalin vs. vehicle) in the expression of various genes in ipsilateral L4-L6 spinal hemicords of male mice 4 days after formalin or vehicle injection. Differentially regulated genes include voltage-activated sodium channels, calcium channels and potassium channels, glutamate receptor channels, sodium/potassium ATPases, cytoskeletal and motor proteins, tetraspanins, and annexins. Red, upregulated, blue, downregulated. Abbreviations: Exc. N., excitatory neurons; Inh. N., inhibitory neurons; Ast., astrocytes; Olig., oligodendrocytes; OPC, oligodendrocyte precursor cells. **(b, c)** Log_2_ fold changes (formalin vs. vehicle) in the expression of genes involved in myelin protein synthesis **(b)**, phospholipid biosynthesis **(c)**, and sphingolipid biosynthesis **(d)**. Abbreviations: *Chka*, Choline kinase alpha; *Cnp*, 2',3'-cyclic nucleotide 3' phosphodiesterase; *Mbp*, Myelin basic protein; *Pcyt1b*, Phosphate cytidylyltransferase 1B, choline; *Sgms1*, Sphingomyelin synthase 1.

*Statistical analysis*: Data were analyzed using unpaired Student’s *t*-tests with false discovery rate (FDR) adjustment. Results are expressed as Log_10_ ion counts and displayed as box-and-whisker plots showing the median and interquartile range. Individual data points represent mice (n=4-7 per group). *P* values reflect comparisons between formalin- and vehicle-treated groups.

**Figure S5: Effects of peripheral injury on the lipidome of purified myelin.** We isolated myelin from bilateral L4-L6 spinal cords 4 days after intraplantar injection of vehicle (gray symbols) or formalin (red symbols) in male mice. Quantification (normalized ion count) of (**a**) cholesterol; (**b-e**) individual species of phosphatidylcholine (PC, **b**), phosphatidylethanolamine (PE, **c**), ceramide (**d**), and sphingomyelin (**e**); (**f**) erucic acid; (**g**) nervonic acid.

*Statistical analyses:* All data were analyzed using unpaired Student’s *t*-tests with false discovery rate (FDR) adjustment and are presented as box-and-whisker plots, where the median and interquartile range are shown. Individual data points represent mice (n = 5-6 per group). *P* values reflect comparisons between formalin- and vehicle-treated groups. ** P < 0.05*; nd, not different.

**Figure S6: Effects of peripheral injury on the lipidome of spinal cord tissue.** We conducted lipidomic analyses on bilateral lumbar spinal cords 4 days after injection of formalin (red symbols) or vehicle (gray symbols) in male mice. Quantification (normalized ion count) of individual lipid species in phosphatidylcholine (PC; **a**), sphingomyelin (**b**), and phosphatidylethanolamine (PE; **c**).

*Statistical analyses:* All data were analyzed using unpaired *t*-tests with false discovery rate (FDR) adjustment and are presented as box-and-whisker plots, where the median and interquartile range are shown. Individual data points represent mice (n = 5-6 per group). *P* values reflect comparisons between formalin- and vehicle-treated groups. ** P < 0.05*; nd, not different.

**Figure S7:** **Imaging mass spectrometry.** Lumbar spinal cords were harvested 4 days after injection of vehicle or formalin in male mice and processed for imaging mass spectrometry. Images shown were reconstructed based on phosphatidylcholine (PC) and phosphatidylethanolamine (PE) as sodium [PC (40:2), PC (36:7)] and potassium adducts [PC (38:3), PE (40:2), PE (38:8), PE (42:10)]. White matter (WM) and gray matter (GM) are highlighted.

**Figure S8:** **Effects of peripheral injury on mean diffusivity in the lumbar spinal cord.** Male mice received injections of vehicle or formalin. Two weeks later, spinal cords were harvested and processed for diffusion tensor imaging. Boxplots show mean diffusivity (mm^2^ per second) in spinal L4 and L5 segments from mice treated with vehicle (gray symbols) or formalin (red symbols).

*Statistical analyses:* Data were analyzed using multiple unpaired Student’s *t*-tests with Bonferroni correction and are presented as box-and-whisker plots, where the median and interquartile range are shown. Individual data points represent mice (n = 5-6 per group). *P* values reflect comparisons between formalin- and vehicle-treated groups.

**Figure S9: Immunofluorescence localization and quantification of BACE1 in the spinal cord.** Lumbar spinal cords were harvested 4 days after injection of vehicle or formalin in male mice and processed for immunofluorescence. **(a)** BACE1 localization in NeuN-positive neurons. Nuclei were stained with 4',6-diamidino-2-phenylindole (DAPI). **(b)** Boxplots showing the number of BACE1-positive nuclei in mice treated with vehicle (gray) or formalin (red).

*Statistical analyses:* Data were analyzed using unpaired *t*-test and are presented as box-and-whisker plots, where the median and interquartile range are shown. Individual data points represent mice (n = 4-6 per group). *P* values reflect comparisons between formalin- and vehicle-treated groups.

**Figure S10: Amyloid plaques in the spinal cord of mice with chronic pain.** (**a**) Fluorescent images of Me-X04-positive amyloid plaques (blue) and IBA-1-positive microglia (red) in the lumbar spinal cords of three separate 8-month-old mice at post-formalin day (PFD) 120. (**b**) Boxplots showing the relative quantification (% area covered) of amyloid-positive deposits in mice treated with vehicle (gray) or formalin (red). Magnification: 40X. (**c**) Fluorescent image of Me-X04 (blue) and IBA-1 immunoreactivity (red) in the lumbar spinal cord of an 11-month-old 5xFAD mouse not exposed to formalin. Magnification: 40X.

**Figure S11: Intrathecal 4G8 administration prevents pain chronification.** Male mice were treated with either formalin or vehicle and, on post-formalin day (PFD) 2 and 4, received intrathecal injections of the Aβ-specific antibody 4G8 (1 μg/μl, 10 μl). Mechanical and heat hypersensitivity and paw edema were assessed at PFD7, PFD14, and PFD21. (**a**) Effects of 4G8 on contralateral mechanical hypersensitivity in animals treated only with vehicle (gray), formalin plus vehicle (red), and formalin plus 4G8 (blue). (**b**) Effects of 4G8 on ipsilateral heat (left) and mechanical (right) hypersensitivity. (**c**) Effects of 4G8 on paw thickness. (**d**) Effects of 4G8 on anxiety-like behavior (elevated plus maze, EPM) including time spent in open arms (seconds), number of closed arm entries, time spent in closed arms, and anxiety index. (**e**) Effects of 4G8 on cognitive performance (novel object recognition test, NOR).

*Statistical analyses*: Two-way repeated-measures ANOVA was used for panels (**a–c**), and one-way ANOVA for panels (**d, e**), followed by the appropriate post hoc analysis. Data are presented as box-and-whisker plots indicating median and interquartile range. Individual data points represent mice (n = 9-10 per group). *P* values reflect comparisons between formalin- and vehicle-treated groups.

**Figure S12: Genetic *Bace1* deletion and pharmacological γ-secretase inhibition block pain chronification.** (**a, b**) Effects of *Bace1* deletion on contralateral (**a**) and ipsilateral (**b**) mechanical hypersensitivity before formalin injection (BL, baseline) and at post-formalin days (PFD) 7 and 14. Symbols: formalin-*Bace1^-/-^* mice (blue), formalin-wild type littermates (red). (**c**) Effects of *Bace1* deletion on ipsilateral heat hypersensitivity before formalin injection (BL) and at PFD7 and PFD14. (**d, e**) Effects of γ-secretase inhibitor NGP555 (10 mg/kg, IP) on ipsilateral (**d**) and contralateral (**e**) heat hypersensitivity measured at PFD7 and PFD14. Symbols: vehicle-vehicle (gray), formalin-vehicle (red), and formalin-4G8 (blue).

*Statistical analyses*: All data were analyzed by two-way repeated measures ANOVA followed by Bonferroni’s multiple comparison. Data are presented as box-and-whisker plots indicating median and interquartile range. Individual data points represent mice (n = 6-9 per group). *P* values reflect comparisons between formalin- and vehicle-treated groups.

**Figure S13: Effects of NAAA inhibition on core energy metabolism and localization of NAAA to oligodendrocytes.** (**a**) Heatmaps showing the effects of NAAA inhibitor ARN19702 (30 mg/kg, IP) administered at post-formalin day (PFD) 2, 3, and 4 in male mice on the expression of genes encoding components of the TCA cycle (**a**) and mitochondrial complexes I, III, and V **(b)**, assessed by bulk RNA-seq at PFD4. Three group of mice (n = 5 per group) were used: vehicle plus vehicle; formalin plus vehicle, and formalin plus ARN19702. Values represent Log_2_ fold changes in gene expression comparing formalin-vehicle vs vehicle-vehicle (FV) or formalin-ARN19702 vs formalin-vehicle (FA). Red, upregulated; blue, downregulated. (**c**) Fluorescent immunostaining of oligodendrocytes (Olig2, green) and NAAA (magenta) in lumbar spinal cord sections. Nuclei were stained with 4′,6-diamidino-2-phenylindole (DAPI, blue). Magnification: ×40. Scale bar, 100 μm.

**Figure S14: Effects of oligodendrocyte *Naaa* deletion on pain chronification.** (**a**) Generation of *Naaa^Olig2-/-^* mice. Crossbreeding of global *Naaa^-/-^* mice with flippase (fl) FLPo-10 mice produced *Naaa^fl/fl^* offspring, which was mated with Olig2-Cre mice to yield the *Naaa^Olig2-/-^* line. Blue boxes: *Naaa* coding sequences; green boxes, Flp recognition target (FRT) sites; green triangles: loxP sites flanking the deleted *Naaa* sequence. The illustration was generated in PowerPoint. Mouse image was from ChemDraw. (**b, c**) Effects of oligodendrocyte-specific *Naaa* deletion on anxiety-like behavior (elevated plus maze): time spent in open arms (left) and number of closed arms entries (right); and (**c**) cognitive function (24-hours novel-object recognition) in mice treated with vehicle or formalin. Red symbols, *Naaa^fl/fl^*; blue symbols, *Naaa^Olig2-/-^.*

*Statistical analyses*: All data were analyzed by two-way repeated measures ANOVA followed by Bonferroni’s multiple comparison. Data are presented as box-and-whisker plots indicating median and interquartile range. Individual data points represent mice (n = 6-9 per group). *P* values reflect comparisons between formalin- and vehicle-treated groups.

**Figure S15: Effects of chronic constriction injury in oligodendrocyte-specific NAAA-null mice.** We subjected male *Naaa^Olig2-/-^* mice and their control littermates (*Naaa^fl/fl^*) to chronic constriction injury (CCI) of the sciatic nerve. Mechanical and heat hypersensitivity were monitored on post-operative days (POD) 14 and 21. Mice were euthanized on POD 28 and lumbar (L4-L6) spinal cord was collected for Aβ quantification. (**a**) Boxplots showing Aβ_42_ (left) and Aβ_40_ (right) levels in insoluble and soluble fractions, respectively, from *Naaa^fl/fl^* (red) and *Naaa^Olig2-/-^* (blue) littermates. (**b**) Effects of oligodendrocyte-specific *Naaa* deletion on ipsilateral and contralateral heat and mechanical hypersensitivity at POD 14 and 21.

*Statistical analyses*: Data were analyzed by unpaired Student’s *t*-test (**a**), or two-way repeated measures ANOVA followed by Bonferroni’s multiple comparison (**b, c**). Data are presented as box-and-whisker plots indicating median and interquartile range. Individual data points represent mice (n = 5-7 per group). *P* values reflect comparisons between formalin- and vehicle-treated groups.

**Reference**

1.	D. Piomelli *et al.*, N-Acylethanolamine Acid Amidase (NAAA): Structure, Function, and Inhibition. *J Med Chem* **63**, 7475-7490 (2020).

2.	Y. Fotio *et al.*, NAAA-regulated lipid signaling governs the transition from acute to chronic pain. *Sci Adv* **7**, eabi8834 (2021).

3.	O. Sasso *et al.*, The N-Acylethanolamine Acid Amidase Inhibitor ARN077 Suppresses Inflammation and Pruritus in a Mouse Model of Allergic Dermatitis. *J Invest Dermatol* **138**, 562-569 (2018).

4.	H. Cai *et al.*, BACE1 is the major beta-secretase for generation of Abeta peptides by neurons. *Nat Neurosci* **4**, 233-234 (2001).

5.	K. Menon *et al.*, The myelin-axolemmal complex: biochemical dissection and the role of galactosphingolipids. *J Neurochem* **87**, 995-1009 (2003).

6.	A. B. Rosenberg *et al.*, Single-cell profiling of the developing mouse brain and spinal cord with split-pool barcoding. *Science* **360**, 176-182 (2018).

7.	T. Kaiser *et al.*, MyelTracer: A Semi-Automated Software for Myelin g-Ratio Quantification. *eNeuro* **8**, (2021).

8.	F. Fiederling, L. A. Hammond, D. Ng, C. Mason, J. Dodd, Tools for efficient analysis of neurons in a 3D reference atlas of whole mouse spinal cord. *Cell Rep Methods* **1**, (2021).

9.	G. J. Bennett, Y. K. Xie, A peripheral mononeuropathy in rat that produces disorders of pain sensation like those seen in man. *Pain* **33**, 87-107 (1988).

10.	Y. Fotio *et al.*, NAAA-regulated lipid signaling in monocytes controls the induction of hyperalgesic priming in mice. *Nat Commun* **15**, 1705 (2024).

11.	A. M. Tagne *et al.*, Metabolic reprogramming in the spinal cord drives the transition to pain chronicity. *bioRxiv*, (2025).
